# Worldwide spread of *Hylurgus ligniperda* (Coleoptera: Scolytinae), and the potential role of bridgehead invasions

**DOI:** 10.1101/2025.05.17.654641

**Authors:** Eckehard G. Brockerhoff, Lea Schläfli, Carolina Cornejo, Julia Kappeler, Jana Orbach, Amira Tiefenbacher, Quirin Kupper, Dimitrios Avtzis, Manuela Branco, Angus J. Carnegie, Kevin D. Chase, Juan Corley, Massimo Faccoli, Elizabeth Gilbride, Brett P. Hurley, Hervé Jactel, Jessica L. Kerr, Natalia I. Kirichenko, Miloš Knížek, Ferenc Lakatos, Victoria Lantschner, Gonzalo Martinez, Nicolas Meurisse, Miguel Angel Poisson, Adrian Poloni, Davide Rassati, Josep M. Riba-Flinch, José P. Ribeiro-Correia, Juan Shi, David Smith, Liam Somers, Yuan Yuan, Simone Prospero

## Abstract

*Hylurgus ligniperda* (F.) is a highly successful invader among bark beetles (Scolytinae) and forest insects in general. Native to the western Palearctic region, it has become established in every continent where its host plants (*Pinus* spp.) occur. Especially in southern hemisphere regions with large pine plantations, it is often highly abundant. As a repeat invader with a wealth of information on various aspects relevant for biological invasions, it is highly suitable as a model organism for studying the role of international trade, the planting of non-native trees, and the potential occurrence of bridgehead invasions (where abundant non-native populations precipitate further invasions). In the present study, our specific objectives were to reconstruct the worldwide invasions of *H. ligniperda* and the pathways involved by using a multi-pronged approach including population genetics, analysis of historic interception data generated from inspections of imports, and records of establishments in the literature. Our review of the native and non-native ranges of *H. ligniperda* and the chronology of establishments revealed at least 13 separate invasions of non-native regions, beginning with Madeira (Portugal) before 1850, and, most recently, eastern China in 2019. We compared the population genetics of 464 specimens from eight countries in the native range and eight countries in the non-native range. Sequencing of the mitochondrial COI gene revealed the presence of 29 haplotypes in six well-supported clades, based on a Bayesian analysis. Non-native populations had significantly lower haplotype diversity (mean *h* = 0.219) than populations in the native range (mean *h* = 0.691). Countries in the non-native range had an average of about two haplotypes compared with about four haplotypes in native countries. In the non-native range, only one or two haplotypes were dominant, and these differed among invaded regions except for haplotype HL-H3 which occurred in Australia, New Zealand and China as well as in four countries in southern Europe, and HL-H4 which was dominant in New Zealand, California, New York State, and eastern China as well as two countries in southern Europe. Analyses of interceptions of *H. ligniperda* with imports arriving in five countries revealed that between 74% and 99% of interceptions originated from other non-native regions while in the USA, most interceptions were linked to imports from the native range beginning in the 1970s. Based on the combined evidence of the chronology of invasions, interception data, and analysis of haplotype distribution, we conclude that the early invasions (before 1950) probably all originated from the native range, while several of the more recent invasions probably originated from parts of the non-native range (suggestive of a bridgehead effect). However, it cannot be determined with certainty what the original sources of each of the invading populations were.

## 1. Introduction

Biological invasions can lead to the decline of native species, threaten the integrity of ecosystems and the provision of ecosystem services, and cause major damage to forests and agricultural crops (Aukema et al. 2010, Kenis et al. 2009, Paini et al. 2016, Vilà and Hulme 2017). New detections of biological invasions continue to occur at a high rate, and there is no sign of a decline or saturation of invaders (Liebhold et al. 2017, Seebens et al. 2018). Bark and ambrosia beetles (Coleoptera: Scolytinae) are particularly successful as invaders of non-native regions, and they include many important forest pests (Hlásny et al. 2021, Lantschner et al. 2020, Vega and Hofstetter 2015). Their invasions are related to a combination of (i) high propagule pressure due to their frequent association with goods transported internationally, (ii) the widespread presence of host plants, (iii) association with plant pathogens, and (iv) the occurrence of inbreeding systems in some species (Brockerhoff et al. 2006a, Brockerhoff et al. 2014, Grégoire et al. 2023, Haack 2006, Vega and Hofstetter 2015). Some invaders are notable for their particularly widespread occurrence. These invasions may either originate from population sources in the native range of species, or because non-native populations spread further from their invaded ranges, so-called secondary or ‘bridgehead’ invasions (Bertelsmeier and Keller 2018, Bertelsmeier et al. 2018, Lombaert et al. 2010). A bridgehead effect may occur especially when non-native populations become highly abundant and potentially more so than in their native region. High abundance of non-native populations may be a result of release from natural enemies (Colautti et al. 2004, Liu and Stiling 2006), enhanced fitness and population growth due to reduced defence of naïve host plants against an antagonist without a shared co-evolutionary history (Desurmont et al. 2011, Gandhi and Herms 2010, Herms and McCullough 2014), or combinations of these and other mechanisms (Brockerhoff and Liebhold 2017, Colautti et al. 2004, Parker et al. 2013). Highly abundant non-native populations may then become an important source, or even the main source, of further invasions into additional regions, potentially causing positive feedback and accelerating invasions (Bertelsmeier and Keller 2018, Bertelsmeier et al. 2018).

The red-haired pine bark beetle, *Hylurgus ligniperda* (F., 1787) (Coleoptera: Scolytinae) is a particularly successful invader. Native to much of Europe, most countries around the Mediterranean Sea, and parts of northern Asia (Western Siberia), it has invaded all continents where its host plants (most if not all species of pine, *Pinus* spp.) occur (Alonso-Zarazaga et al. 2023, Brockerhoff et al. 2014, Lin et al. 2021, Seybold et al. 2016, Wood and Bright 1992). The native and non-native ranges of *H. ligniperda* have been listed and reviewed by numerous authors, with some disagreement. In part, this is due to several incorrect occurrences mentioned in the literature which have been propagated in later publications. For example, Wood and Bright (1992) list its presence as a native species in Norway, China, and Japan, all apparently incorrectly. The uncertainty about the presence and status of *H. ligniperda* in north-east Asia (CABI 2021, Lin et al. 2021, Wood and Bright 1992) has been resolved (Lin et al. 2021), but questions remain about several other regions. Nevertheless, there are no questions about the status of *H. ligniperda* (i.e., its presence as a non-native established species) in most of its non-native range and about its very high abundance in many invaded regions, especially in parts of the southern hemisphere where large pine plantations occur (Brockerhoff et al. 2006b, Brockerhoff et al. 2025, Faccoli et al. 2020, Lantschner et al. 2020). This multitude of invasions makes *H. ligniperda* a suitable case study to reconstruct origins and pathways of invasions. Specifically, the sequence of repeated invasions over the course of more than 150 years and the high abundance in many non-native regions suggest that some of these invasions may be the result of secondary or ‘bridgehead’ invasions.

Testing the hypothesis that invasive populations can become an important source of further invasions (via the ‘bridgehead’ effect) can advance our understanding of processes driving biological invasions. However, it is difficult to determine the source of invasive insect populations because most such invasions are accidental and typically go unnoticed for many years until they are discovered eventually. The rarity of observations of insect invasions is due to a lag phase (between the first establishment and the detection of an invasive population) which is well-known for plant invasions (e.g., Duncan 2021, Robeck et al. 2024) but rarely shown in insect invasions (Siegert et al. 2014). This lag makes it very difficult to trace back the source of invading populations and the exact mechanism of transport to a non-native region. Data on border interceptions of non-native insects, which are recorded by customs officers carrying out phytosanitary inspections of imports (Brockerhoff et al. 2014, Turner et al. 2025), can shed some light on the potential sources of such organisms (Brockerhoff et al. 2003, Brockerhoff et al. 2014, McCullough et al. 2006, Turner et al. 2021, Worm et al. 2024).

Another approach to determine the possible origin of non-native populations is to compare their genetic background with that of native (and other non-native) populations and look for similarities. The mitochondrial gene Cytochrome Oxidase I (COI) is a widely used molecular marker in phylogenetic studies (Cognato and Sperling 2000, Víctor and Zúñiga 2016) and population genetic studies of insects (Dong et al. 2021), including Scolytinae and other beetles (e.g., Fernandez et al. 2021, Ito et al. 2008, Mayer et al. 2015). It is suitable for these purposes due to its near-neutrality, lack of recombination, and a clock-like evolutionary rate. Although nuclear microsatellites may provide a higher resolution than COI (Taerum et al. 2016, Tsykun et al. 2019), their use with Scolytinae has not always been successful for distinguishing regional populations because their spatial structure may be limited (Gugerli et al. 2008). Genomic tools such as RAD sequencing are used increasingly for population genetic studies; however, the results from such studies of Scolytinae were largely consistent with those from using COI (Urvois et al. 2023). Combining both approaches, analyses of population genetics together with border interception data should provide more insights about long-distance, human-assisted movements of insect populations.

In the present study, our objectives were:

1. To review the worldwide presence, native and non-native status, and first records of invasions (i.e., confirmation of establishment of non-native populations) of *H. ligniperda*;
2. to analyse the genetic structure of populations by sequencing the mitochondrial COI gene;
3. to compile and review sources of border interceptions of *H. ligniperda* in multiple countries associated with imports from worldwide sources;
4. and to use the combined findings to obtain plausible scenarios of native or non-native sources of each invasion of *H. ligniperda*.

## 2. Materials and Methods

### 2.1 Reconstruction of the worldwide presence, native and non-native status, and first records of invasions of *H. ligniperda*

Information about the presence of *H. ligniperda* and its status was compiled from multiple searches for “ligniperda” AND “Scolyti*” of the ‘Web of Science Core Collection’ (WOSCC, all fields, last on 15/03/2025) and Scopus (all fields, last on 15/03/2025). These searches revealed 59 records in WOSCC (the oldest being Neumann 1979) and 228 documents in Scopus (the oldest being Elton et al. 1964). Therefore, additional and especially older references were searched for in monographs and review articles on Scolytinae including Alonso-Zarazaga et al. (2023), Balachowsky (1949), CABI (2021), Eichhoff (1881), Pfeffer (1995), Wood and Bright (1992), and others mentioned below, and by contacting colleagues in all regions of interest (e.g., Argentina, Australia, the Azores, Czechia, Japan, China, Madeira, Russia, South Africa, the USA) for further information, literature, verification, and, where relevant, translations to English. The earliest publication found by back-searches that mentioned an invasion of *H. ligniperda* was Wollaston (1854). The results of these searches were synthesised and tabulated.

### 2.2 Population genetic analyses

#### 2.2.1 Sampling

Specimens of *H. ligniperda* used in this study originated from 16 countries on six continents (**Table 1**, **Suppl. Table 1**). Some samples originated from the study by Faccoli et al. (2020). To have better coverage across the distribution range of *H. ligniperda,* numerous additional samples were collected between 2016 and 2022 in eight countries in the native range (Czechia, France, Greece, Hungary, Italy, Portugal, Spain, Switzerland) and nine countries or states in non-native ranges (Argentina, Australia, Chile, China, New Zealand, South Africa, Uruguay, and the USA (California and New York State)) (**Suppl. Table 1**). Unsuccessful attempts to collect *H. ligniperda* occurred in Croatia, Germany, Romania, Siberia (near Krasnoyarsk), and Sweden (but note that this does not mean that the species is not present there). In addition, several specimens of *Hylurgus micklitzi* Wachtl, 1881 were obtained from Portugal and Greece and used for comparison.

**Table 1.**
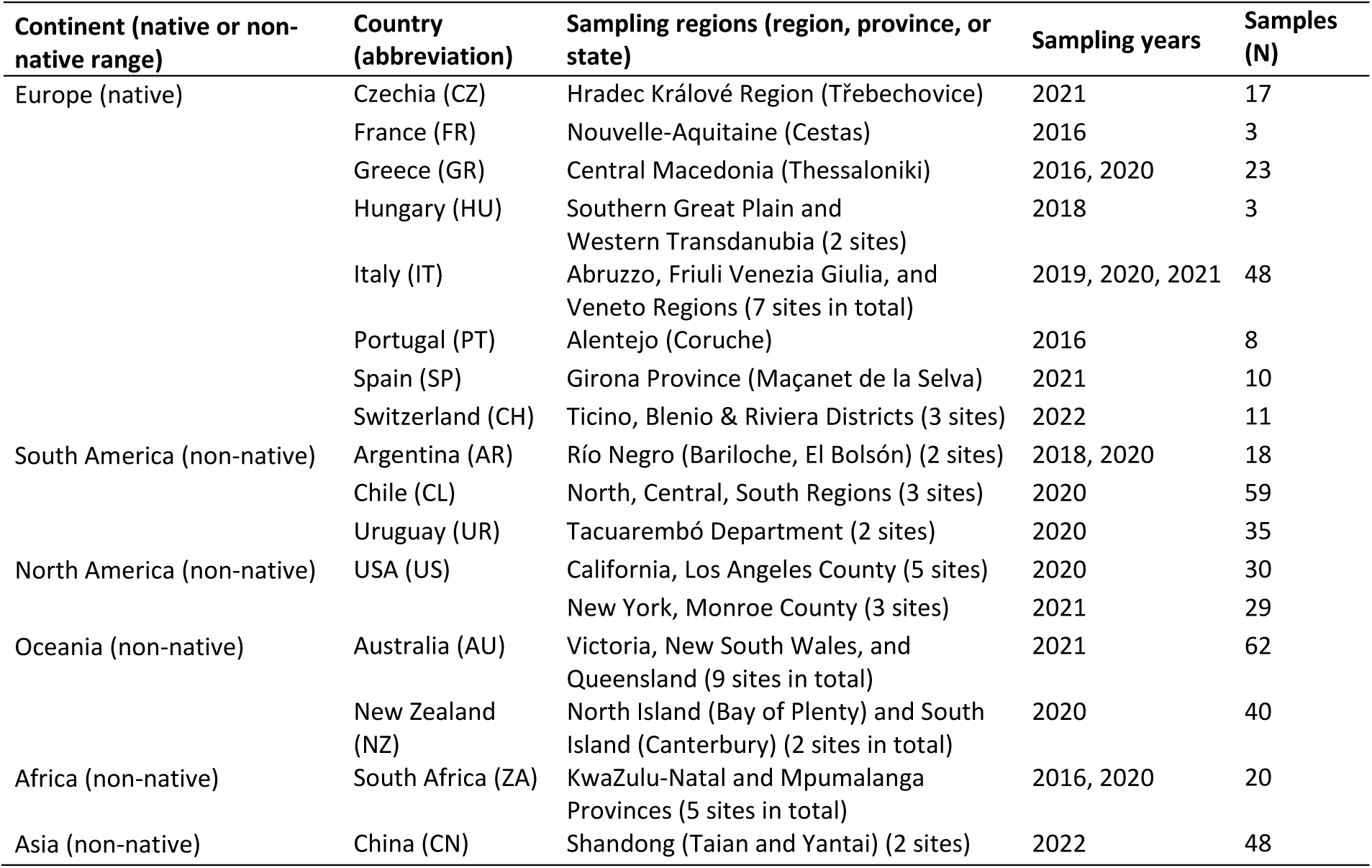
Geographic origin of the 464 *Hylurgus ligniperda* specimens analysed in this study (for further details about the sampling locations see Suppl. Table 1)

Specimens of *H. ligniperda* were identified based on morphological features using the identification key of Grüne (1979) and by comparison with reference specimens. Specimens were stored in 95% ethanol and sent to the Swiss Federal Institute for Forest Snow and Landscape Research WSL for molecular analyses. DNA was extracted from 508 out of the 1640 specimens collected. More than two thirds of specimens were not used for genetic analyses because for some locations we obtained more samples than required. From 464 specimens, good-quality DNA was obtained that could be used for COI sequencing (**Table 1**). The correct identity of samples from the *H. ligniperda* haplotypes we identified (see Results) was verified by comparing their COI sequences with those in the public animal library in BOLDSYSTEMS (https://id.boldsystems.org/) (Ratnasingham et al. 2024) (see below for further details).

#### 2.2.2 DNA extraction, PCR amplification and sequencing

For DNA extraction, up to four legs were removed from each beetle with sterile tweezers under a microscope and transferred into a 1.5 ml microtube. After lyophilization, the legs were ground with a sterile pestle directly in the microtube. The DNA was subsequently extracted using the NucleoSpin Tissue XS Kit (Macherey-Nagel, Düren, Germany) according to the manufacturer’s instructions.

PCR amplification of COI was performed using the primer pair C1-J-1718 (Simon et al. 1994) and CO1-a2411 (Normark et al. 1999) in a reaction volume of 30 μl containing 15 μl JumpStart REDTaq ReadyMix (Sigma Aldrich: Merck KGaA, Darmstadt, Germany), 3 μl DNA template, 9.6 μl molecular grade water (Merck) and 1.2 μl of each primer (10 μM). The thermal cycling parameters were set as follows: 94°C 2 min initial denaturation, five cycles at 94°C 20 sec, 47°C 20 sec, and 72°C 30 sec, followed by 40 cycles at 94°C 20 sec, 52°C 20 sec, and 72°C 30 sec and a final extension of 10 min at 72°C. The PCR amplification was verified on 1.5% agarose gel. Successfully amplified products were enzymatically purified by exonuclease I and alkaline phosphatase treatment according to the manufacturer’s instructions (GE Healthcare, Chicago, Illinois, USA) and Sanger sequenced on an ABI PRISM 3130 Genetic Analyzer (Applied Biosystems) using the same primer pair as for the PCR reaction. Sequencing reactions were conducted in 7 μl mixtures, using the Big Dye Terminator 3.1 cycle sequencing premix kit (Applied Biosystems: ThermoFisherScientific, Waltham, Massachusetts, USA). Forward and reverse raw sequences were trimmed and assembled in CLC Genomic Workbench 20 version 20.0.4 (CLC bio: QIAGEN, Hilden, Germany). Forty-four out of 508 COI sequences from *H. ligniperda* we obtained were excluded from further analyses because of insufficient quality. The final sequence alignment consisted of the quality-checked sequences with a length of 692 bp and no gaps. A representative sequence of each of the 29 haplotypes was made publicly available in GenBank (PV384201-PV384229).

#### 2.2.3 Haplotypes and genetic diversity

The genetic diversity across the whole dataset and within each geographic population (by country or US state) was estimated using the number of observed haplotypes, haplotype diversity (*h*), nucleotide diversity (*π*) and Tajima’s D statistic (Tajima 1989). These analyses were conducted in the R (version 4.2., R core Team 2022) environment Rstudio (RStudio Team 2022) using custom scripts. DNA was read in with the package ape (version 5.7-1, Paradis and Schliep 2019), and haplotypes were generated using the function *haplotype* from the package pegas (version 1.2, Paradis 2010). The *h* and *π* values were calculated for the different populations or across the dataset using functions *hap.*div, specifying the method ‘Nei’ and *nuc.div*, and values for Tajima’s D test statistic and corresponding *P*-values (beta) were calculated with the function *tajima.test* from pegas, respectively. Mean haplotype diversity in the native and non-native ranges was compared with a weighted t-test, after testing for normal distribution using the Shapiro-Wilk test. The square root of the sample size of each sample was used as the weight variable in the t-test.

The genetic relationship between the different COI haplotypes was further explored and visualised by constructing a haplotype network using the function *haploNet* based on a median-joining algorithm (Bandelt et al. 1999) from pegas. For an overview of the distribution of the different haplotypes across countries and populations, frequencies of the haplotypes were summarised across populations, and these were plotted using R packages ggplot2 (version 3.4.2, Wickham 2016) and scatterpie (version 0.1.8, Yu 2024).

#### 2.2.4 Phylogenetic analysis

To identify identical sequences, the alignment containing 464 samples was analyzed in SplitsTree5 (Huson et al. 2006a, 2006b) using the “Group Identical Haplotypes” option, which collapses identical sequences into a single sample. The phylogenetic relationship of the 29 distinct haplotypes identified in SplitsTree5 (see Results) was explored with Bayesian statistics using BEAST v1.10.4 (Suchard et al. 2018) on the CIPRES Science Gateway (Miller et al. 2010). The Smart Model Selection in PhyML, SMS v.2.0 (Lefort et al. 2017) and the Akaike Information Criterion (AIC) (Akaike 1973) were used as implemented on the ATGC Montpellier bioinformatic platform (http://www.atgc-montpellier.fr) to select the model that best fitted the data. BEAST analysis was inferred with the substitution model HKY85 (Hasegawa et al. 1985) and using the BEAGLE library v3.1.2 (Ayres et al. 2012) for accelerated, parallel likelihood evaluation. The analysis was run with 160 million states, sampling every 1,000th state and ignoring the first 20 million states as burn-in. The convergence and the consequent proportion of burn-in were assessed using the desktop software Tracer v1.7.2 (Rambaut et al. 2018) included in the BEAST software package. To obtain the Bayesian posterior probabilities (PP), a maximum clade credibility tree was generated by analysing the sampled trees produced by BEAST in TreeAnnotator v.1.10.4 (BEAST software package). The best tree including the posterior probabilities of the nodes was represented as unrooted phylogram using TreeGraph v2 (Stöver and Müller 2010).

### 2.3 Interception data

Data on interceptions of *H. ligniperda* and other Scolytinae that were found during inspections of goods imported from origins around the world were obtained from national plant protection organisations of several countries and from published datasets: Australia (covering the period 2003-2023), Chile (1995-2005), Japan (2002-2019), New Zealand (1950-2024), and the USA (1914-2024). For selected datasets, we had additional data on the country or region of origin and the nature of commodities and associated packaging of individual interception events (e.g., New Zealand 1950-2000 and USA 1914-2008). Note that interception data represent ‘interception events’ whereby each event may have included a single or many individuals of a species. For more information on the sources interceptions and inspection procedures that generated these data, see Brockerhoff et al. (2006a), Brockerhoff et al. (2014), Haack (2006), Ide et al. (2014), and Turner et al. (2025).

Origins of interceptions and native or non-native status of origins were grouped by region whereby Europe, North Africa and West Asia (e.g. Turkey) were considered native and East Asia, Oceania, North America, South America and Southern Africa were considered non-native. Cases of interceptions of *H. ligniperda* where origins were missing as well as origins from countries where *H. ligniperda* is not known to occur (Canada, Hong Kong, India, Indonesia, Solomon Islands, Taiwan, and Vietnam) were classified as ‘unknown’ origins. Such cases may be due to reuse of wood packaging materials for subsequent shipments. Because populations of *H. ligniperda* in the USA and China established only relatively recently (detections in 2000 and 2019, respectively) and are still restricted to smaller areas of limited extent, interceptions with shipments from these countries were considered less likely to be correct and also classified as ‘unknown’ origins.

## 3. Results

### 3.1 Reconstruction of the worldwide presence, native and non-native status, and first records of invasions of *H. ligniperda*

**Native range:** Our synthesis of records of occurrences and absences is given in **Suppl. Table 2** and summarised in **Table 2**. For Europe, North Africa and Asia, we mostly agree with the assessments of Alonso-Zarazaga et al. (2023, 2025) and Bright (2021), but several updates are needed (see below).

**Table 2.**
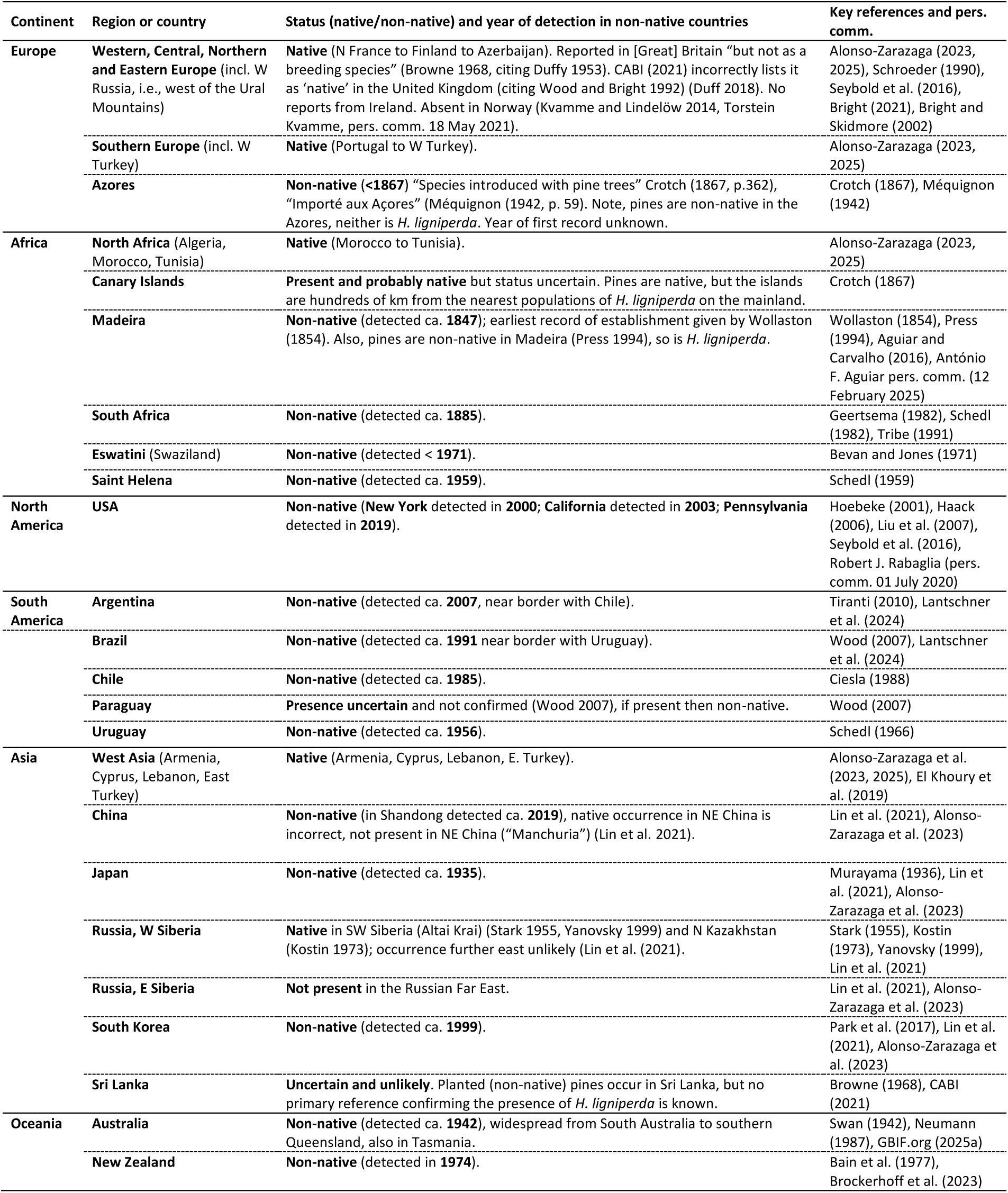
Synthesis of occurrence of *H. ligniperda* as a native or non-native species by continent and country or region based on a major review of the literature, collection of specimens and correspondence with experts. The year of first detection or reporting as established in non-native regions is given. Information compiled from a review of Balachowsky (1949), Browne (1968), Wood and Bright (1992), Pfeffer (1995), Seybold et al. (2016), Alonso-Zarazaga et al. (2023), CABI (2021), Lin et al. (2021), as well as key references cited in the right column. For details see Suppl. Table 2.

There is widespread agreement in the literature about the native range of *H. ligniperda* in much of temperate and southern Europe. However, there is some uncertainty about its presence in northwest and northern Europe, and in the Azores (see below). According to Duffy (1953) and Browne (1968), it is reported from [Great] Britain but “not as a breeding species”, while Duff (2018) lists *H. ligniperda* under “non-established introductions”. There are no records of occurrence in the UK or Ireland in GBIF.org (2025a, 2025b); however, specimens have been trapped in southeast England (Daegan Inward, pers. comm. 31 March 2025). In Scandinavia, there are records from Denmark (Alonso-Zarazaga et al. 2023, 2025), but it is absent from Norway (Kvamme and Lindelöw 2014). It has been recorded occasionally in southern Sweden (Åke Lindelöw, pers. comm. 18 May 2021), and further north along the Baltic coast of Sweden (GBIF.org 2025a, 2025b), as well as in Finland and the Baltic countries (Alonso-Zarazaga et al. 2023, 2025, Bright 2021). In the European part of Russia, *H. ligniperda* is present but apparently not in the colder, boreal areas. According to Stark (1955), it occurs in Smolensk Oblast, Voronezh Oblast, and in the Caucasus region (which appears to extend into Georgia and Azerbaijan). Stark (1955) also mentions its occurrence in Belarus and Ukraine including Crimea. Alonso-Zarazaga et al. (2023, 2025) list Georgia and Azerbaijan as part of its native range.

Although some consider *H. ligniperda* to be mainly a Mediterranean species, there are many records in GBIF from Belgium, the Netherlands, Germany, Austria, Poland, Czechia, and Slovakia (GBIF.org 2025a, 2025b). Already in the late 1800s, Eichhoff (1881) summarised its native range as including Central and Southern Europe and that it is present “in all of Germany, France and Austria”. However, based on our review of the wider literature, it is generally most abundant in southern Europe from Portugal to Greece and western Turkey, especially in areas where pines are abundant.

In North Africa, *H. ligniperda* is present from Morocco to Tunisia and in the Canary Islands. In the Canary Islands, a native species of pine (*Pinus canariensis*) occurs, and *H. ligniperda* is probably also native, despite the distance of the Canary Islands from the African mainland. We are unaware of any records from Libya to Egypt. In Western Asia, it occurs in Cyprus, Lebanon and Turkey, and probably also in the coastal part of Syria between Turkey and Lebanon, but it appears to be absent from Israel (Mendel 1987). In Central Asia, *H. ligniperda* has been recorded (and is assumed to be native) in Altai Krai (in the southern part of Western Siberia) (Stark 1955, Yanovsky 1999, Alex Petrov, pers. comm. via Natalia Kirichenko 17 February 2025) and in adjacent areas of northeast Kazakhstan (Kostin 1973). It is not known if this is an isolated population or if the species occurs continuously from Altai Krai westwards along the southern border of Russia.

### Non-native range

The first known establishments of *H. ligniperda* as a non-native species were in Madeira and in the Azores Archipelagos. Although many if not all publications since the 1990s which summarised the distribution of *H. ligniperda* considered Madeira to be part of the native range (e.g., Alonso-Zarazaga et al. 2023, CABI 2021, El Khoury et al. 2019, Wood and Bright 1992), it is clearly not native **in Madeira**. Wollaston (1854) provided the first record of its occurrence in 1847 in cultivated (non-native) pine forests where it is (like *Tomicus piniperda*) “in all probability […] naturalized in Madeira”. Balachowsky (1949) also considered it ‘introduced’ in Madeira. Given the narrow host range of *H. ligniperda* and that neither pines nor other Pinaceae are native in Madeira (Press 1994), there can be no doubt that it is a non-native species there (Aguiar and Carvalho 2016, António F. Aguiar, pers. comm. 12 February 2025) which was probably introduced with pine timber. The same situation, with both pines and *H. ligniperda* being non-native species, applies to the population in the **Azores** where an invasion occurred before 1867 (Crotch 1867, Méquignon 1942).

The next recorded invasion of *H. ligniperda* appears to be in **South Africa** where it was detected in 1885 (Geertsema 1982, Schedl 1982, Tribe 1991). Some recent studies (e.g., Lantschner 2024) refer to 1973 as the year of detection in South Africa, but it had been detected there nearly 90 years earlier. Schedl (1959) reported the presence of *H. ligniperda* in Saint Helena (in the Atlantic off the coast of Angola). The occurrence of *H. lig*niperda in Eswatini (formerly Swaziland) was reported by Bevan and Jones (1971), but it was likely present many years before that given that Eswatini is partially surrounded by South Africa.

For **northeast Asia**, there was some disagreement whether *H. ligniperda* is native or non-native in this area. Wood and Bright (1992) stated it is present in “Manchuria in China” and Japan without indicating that it is introduced (i.e. it was considered native). Its potential occurrence in northeast Asia was reviewed by Lin et al. (2021), and it was concluded that it is not native anywhere in northeast Asia. The first detection of a non-native population in Asia was in **Japan** in 1935 (Murayama 1936, Lin et al. 2021). In **South Korea**, the earliest recorded specimens are from 1999, and the studies by Park et al. (2017) and Lin et al. (2021) confirmed that it is clearly established and widespread. In **China**, the presence of a non-native population was confirmed in Shandong Province in 2019 (Lin et al. 2021), although it probably became established there years before 2019 as it was found already over a larger area. In the Asian part of Russia (i.e., east of the Ural Mountains), the situation was also unclear. This was also reviewed by Lin et al. (2021) who stated “… this beetle is not found in eastern Russia (Alex Petrov, personal communication)”. However, as *H. ligniperda* is present in Western Siberia in Altai Krai (Stark 1955, Yanovsky 1999) and in northwest Kazakhstan (Kostin 1973), we consulted with forest entomologists in Siberia and Moscow (Alex Petrov, pers. comm. via Natalia Kirichenko 17 Feb. 2025). They confirmed the records of *H. ligniperda* from Altai Krai and northwest Kazakhstan (Leninogorsk Forest), and that there are indeed no records of a population in the Russian Far East. Browne (1968) listed its occurrence in **Ceylon (i.e. Sri Lanka)** without giving a source. Browne (1968) was probably the source of subsequent mentions of occurrence in Sri Lanka (Bain 1977, Brockerhoff et al. 2003, Seybold et al. 2016, CABI 2021), but we are not aware of any primary reference or recent publications that confirm its presence there. There are plantations of *Pinus caribaea* in Sri Lanka (Ashton et al. 1997) but given the climate and lack of any known primary references of occurrence, we assume that *H. ligniperda* is not present in Sri Lanka. It is possible that specimens intercepted with imports could have led to the assumption that it occurs in Sri Lanka.

In **Australasia and Oceania**, the first record of an established population was in 1942 in southernmost South **Australia** (Swan 1942). Today, it is present also in Victoria, Tasmania, New South Wales and southern Queensland. In **New Zealand**, a population was first detected in 1974 in Whitford near Auckland (Bain 1977, Brockerhoff et al. 2003) from where it has spread across most of the North Island and South Island.

In South America, the first established population was detected **in Uruguay** in 1956 (Schedl 1966) where afforestation with pines began in the 1930. A disjunct population was detected in **Chile** in 1985 (Ciesla 1988) and on the eastern side of the Andes in **Argentina** in 2007 (Tiranti 2010). The species is also present in **Brazil**. Wood (2007) gives the first record in Brazil as 1981 with reference to a paper by Schönherr and Pedrosa-Macedo (1981), although that paper only mentions the presence of *H. ligniperda* in Uruguay (in Punta del Este which is about 200 km from the nearest border with Brazil). Another record from Brazil in Wood (2007) is from 1991 from an ethanol-baited trap in Rio Grande do Sul State (which shares its southern border with Uruguay). Lantschner et al. (2024) also consider 1991 the first record for Brazil. A second invasion affecting pine plantations in northeastern Argentina near the borders with Brazil or Uruguay occurred later. The presence of *H. ligniperda* in Paraguay is uncertain (Wood 2007, Lantschner et al. 2024), but given the occurrence of pine plantations near plantations in Brazil and Argentina, its presence there is not implausible.

In the **USA**, the first breeding population was detected in 2000 in northern **New York State** near Rochester after two specimens had already been trapped nearby in 1994 and 1995 (Hoebeke 2001). Subsequently, specimens have been trapped at other sites in Monroe, Niagara, Broome and Jefferson counties, and in 2019 also in Pennsylvania in Crawford County (Robert J. Rabaglia, pers. comm. 1 July 2020, Seybold et al. 2016). In 2003, a population was detected in southern **California** (Haack 2006, Liu et al. 2007, Seybold et al. 2016). Based on records until 2020, *H. ligniperda* appears to be restricted to southern California in Kern, Los Angeles, San Diego, San Luis Obispo and San Bernardino counties.

We conclude that there were **at least 13 independent invasions** of geographically separated regions across all continents where pines occur as native or non-native plants (**Table 2**). Several records in the literature need to be updated with the information we provide above.

### 3.2 Haplotype diversity

The final data set contained 464 sequences with a length of 692 nucleotides and 41 polymorphic sites. Grouping of these sequences into identical sequences resulted in 29 different haplotypes (**Table 3**). Checks for species identity of all these haplotypes with sequences in BOLDSYSTEMS (https://id.boldsystems.org) (Ratnasingham et al. 2024) confirmed in all cases the species *Hylurgus ligniperda* with 98-100% identity. These were all clearly distinguished from *Hylurgus micklitzi* and *Hylurgus indicus* Wood, the only two other species in this genus.

**Table 3.**
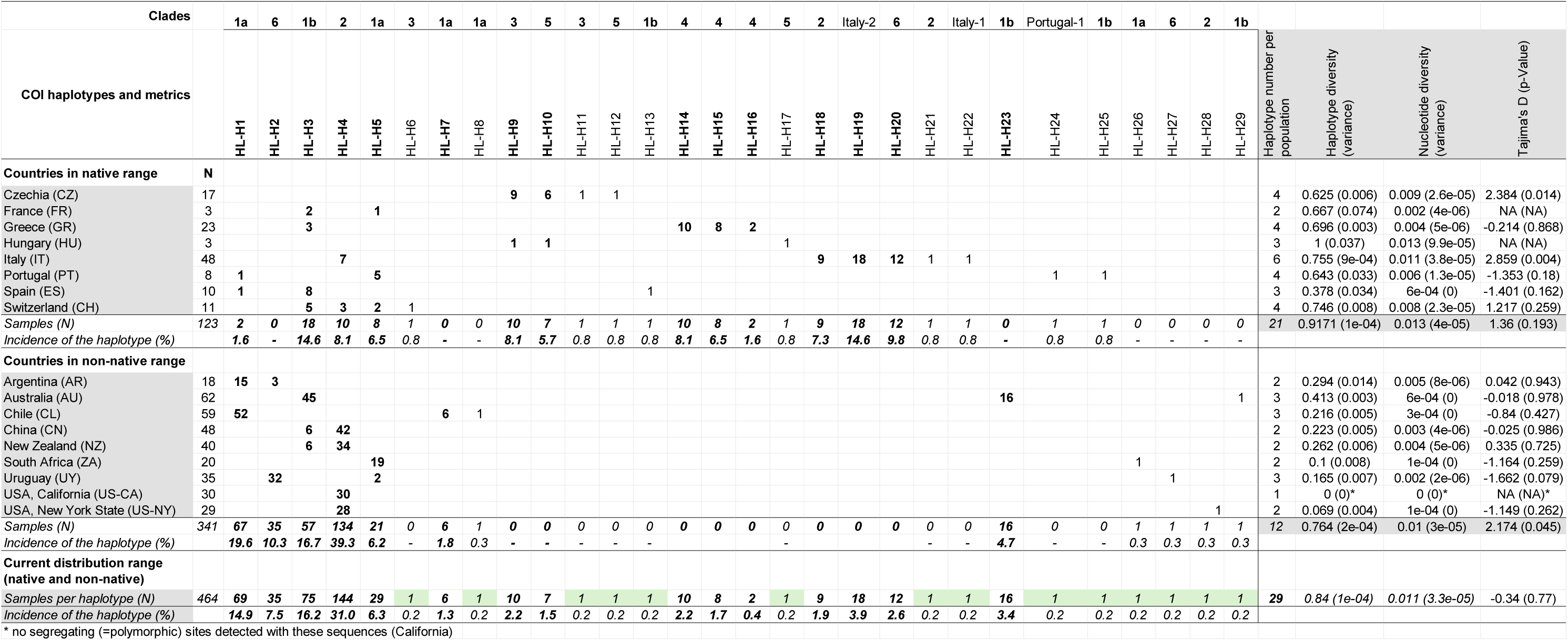
Overview of COI haplotypes of *H. ligniperda* across the entire dataset. Haplotypes which were found in more than one sample are marked in bold font (N=15), while 14 of the 29 total haplotypes were only found once. For haplotype diversity *h* and nucleotide diversity *π*, variances are indicated in brackets and for Tajima’s D, the P-value is indicated in brackets.

Of the total of 29 COI haplotypes identified in the overall *H. ligniperda* population, 14 consisted of only one sample while the others included between two (HL-H16) and 144 (HL-H4) samples (**Table 3**). The highest haplotype diversity was observed in the native range of the species in Europe (in total: 21 haplotypes, overall *h* = 0.9171 ± 1e-04), whereby samples from individual countries were composed of two to six haplotypes (*h* = 0.667 ± 0.074 and 0.755 ± 9e-04 in France and Italy, respectively) (**Table 3**). Despite the more or less continuous occurrence across most of the native range, the majority of haplotypes were found in only one or two countries in the native range, and only two of them occurred more widely (i.e., HL-H5 in three countries and HL-H3 in four countries).

In the non-native range, only 12 haplotypes were identified, with HL-H4 being the most common (39.3% of non-native samples), even though the non-native range was sampled nearly three times more intensively and, collectively over a larger geographic range, than the native range (341 vs. 123 samples, respectively). Noteworthy, eight haplotypes from the non-native range (HL-H2, HL-H7, HL-H8, HL-H23, HL-H26, HL-H27, HL-H28, HL-H29; **Table 3**) were not detected in the native range. The haplotype diversity (*h*) in non-native populations ranged from 0 (in California, one single haplotype detected) to 0.413 ± 0.003 (in Australia) (**Table 3**). Overall *h* across all non-native populations was 0.764 (± 2e-04). Mean haplotype diversity in the non-native range (mean *h* = 0.219) was significantly lower than in the native range (mean *h* = 0.691) (two-sample weighted t-test, P < 0.001). Overall nucleotide diversity (*π*) was only slightly lower in the non-native range (0.010 ± 3e-05) than in the native range (0.013 ± 4e-05) (**Table 3**). Tajima’s D across all non-native populations was 2.17 and statistically significant (P = 0.045), which indicates that rare alleles are scarce, although for individual populations (countries), not a single D-value was statistically significant (**Table 3**). In the native range overall, D was 1.36 and appeared to be lower (i.e., more rare alleles), compared with the non-native range, but this was not statistically significant (P = 0.193). However, in some individual countries in the native range, D was relatively large and significant [Italy, D = 2.86 (P = 0.004) and Czechia, D = 2.38 (P = 0.014)], while in other countries, it was negative, although not significant).

The map of haplotypes (**Figure 1**) clearly illustrates these patterns of a much higher haplotype diversity in the main part of the native range of *H. ligniperda* in Europe than in the non-native ranges. In the non-native range, few haplotypes are present, and in all cases only one or two were found to be dominant. For example, HL-H1 dominates in Chile and Argentina, HL-H2 in Uruguay, HL-H3 and HL-H23 in Australia, HL-H4 in New Zealand, California, New York, and China, and HL-H5 in South Africa (**Table 3**, **Figure 1**). Haplotype HL-H4, which is particularly widespread and common in the non-native region, is also among the more common haplotypes in the native range (in Italy and Switzerland). Interestingly, the other haplotype that is well represented in non-native regions, HL-H3 with three occurrences in Australia, New Zealand and China, occurs also in multiple countries in the native range including France, Greece, Spain and Switzerland.

**Figure 1.**
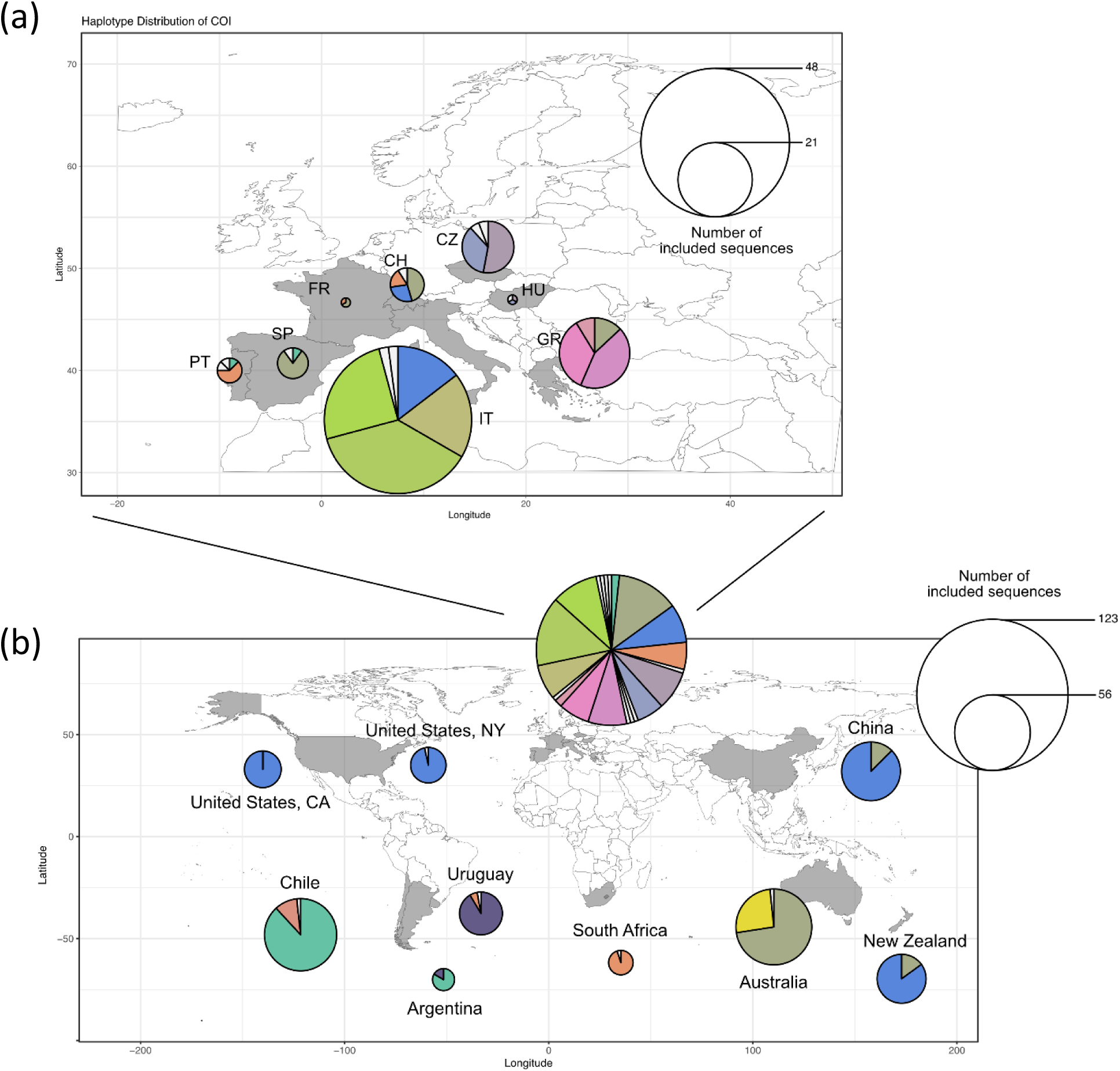
Maps showing the distribution of COI haplotypes of *Hylurgus ligniperda* examined in the present study which originated from (a) the native range in Europe, and (b) the invaded regions in other continents. The area of segments is proportional to the number of samples representing particular haplotypes (as displayed in the circle legend). In total, N = 29 haplotypes were identified. Six unique haplotypes in the native range (which were identified only once and in only one country in the native range) are coloured in white.

### 3.3 Phylogenetic analysis and haplotype network

The Bayesian analysis we performed with the 41 unique patterns from 692 nucleotide sites resulted in a best tree (from 162,615 trees sampled) which is presented as an unrooted phylogram in **Figure 2**. Including the split of clade 1 into (sub-)clades 1a and 1b, seven well-supported clades were identified, and three specimens (Italy-1, Italy-2 and Portugal-1) remained as unique, distinct haplotypes. Clades 1a, 1b, 2 and 6 contained specimens of European and non-European origin, all others only European specimens. The four clades which are present in the non-native range are partly separated between regions. For example, clade 6 is present only in Uruguay and Argentina in the non-native range, and the two parts of clade 1a occur only in Uruguay and South Africa or in Chile and Argentina. However, clade 1b is present in Australia, New Zealand and China, while clade 2 is present in New Zealand, California, New York and China.

**Figure 2.**
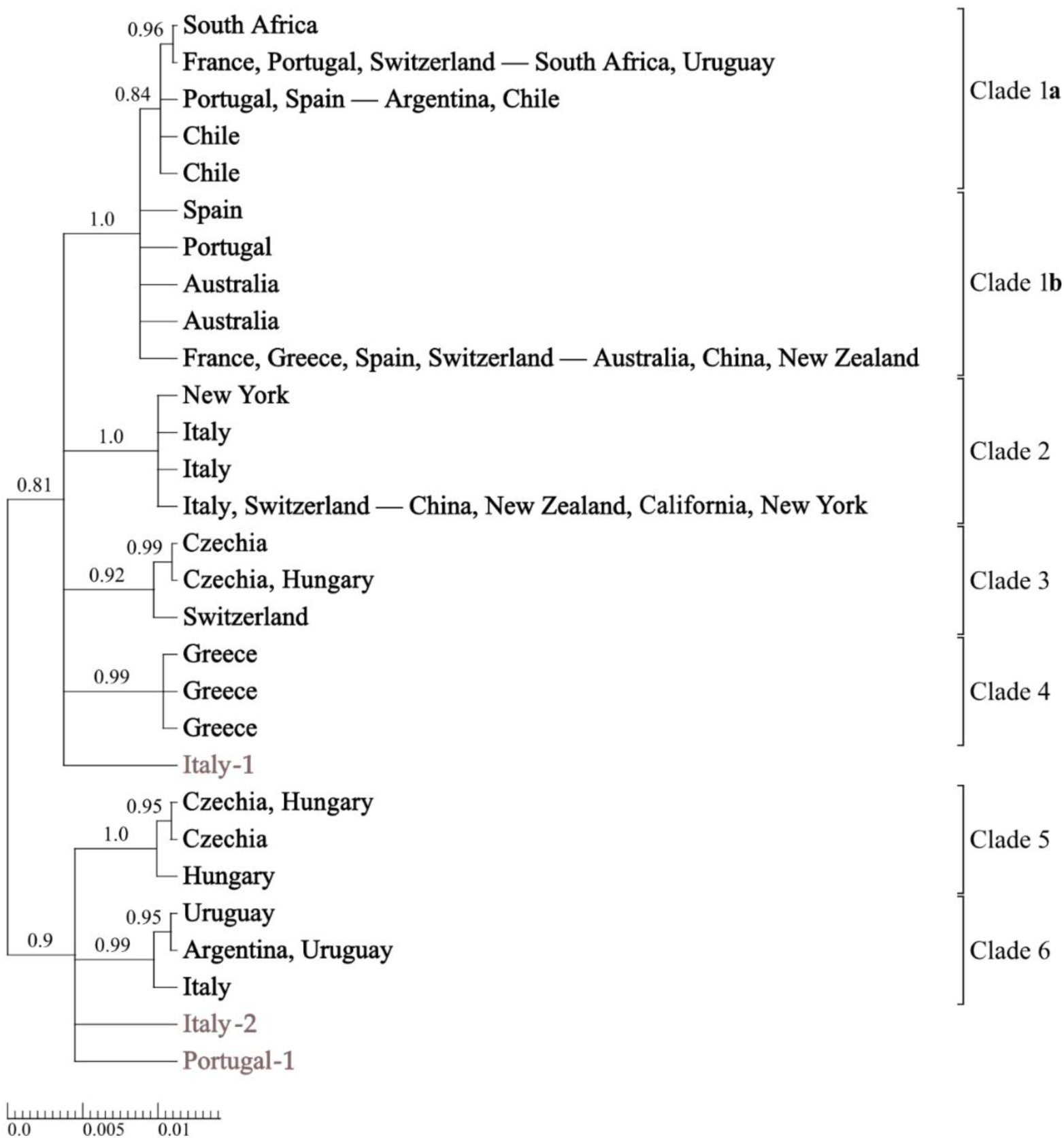
Unrooted Bayesian best tree of *Hylurgus ligniperda* based on 29 different COI sequences and labelled with the geographic origin of all samples with a common haplotype. The long hyphen separates European from non-European specimens. Branches with posterior probabilities PP<0.8 were collapsed, all other posterior probabilities are plotted above the branches.

The clades identified by the Bayesian analysis were consistent with the haplotype network (**Figure 3**). All clades but clade 4 were clearly dominated by a single haplotype, which included 65-93% of the clade samples. Clade 1, subdivided into two sub-clades, was the most diverse one, with a total of 10 haplotypes which differed by maximum two polymorphic sites (within sub-clade). The four haplotypes within clade 2, among them the most common HL-H4 (144 samples), were also genetically closely related. The other clades (clade 6 and especially clades 3, 4, and 5) each included three less frequent haplotypes, with a larger within-clade distance among haplotypes compared to clades 1 and 2.

**Figure 3.**
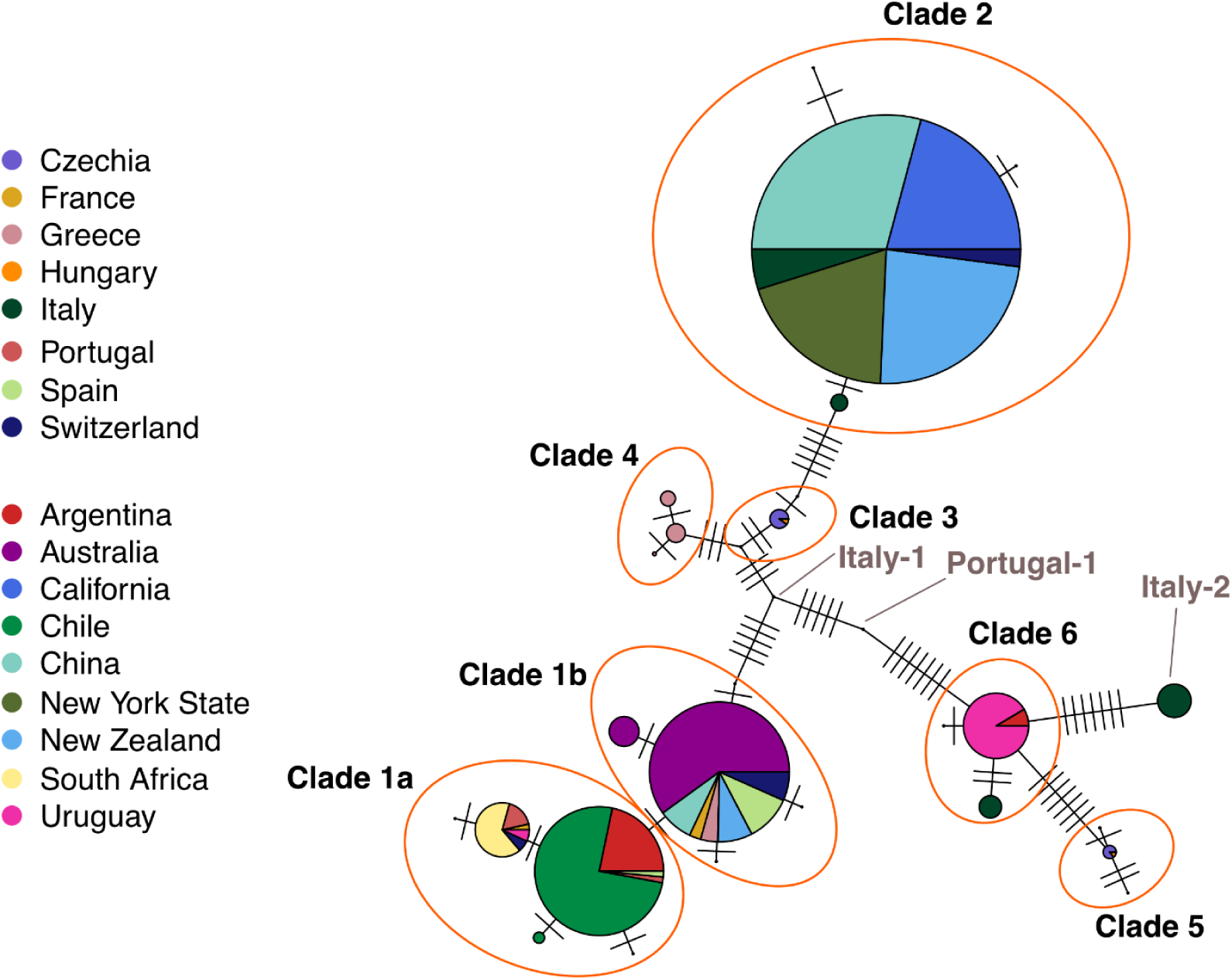
Network of the 29 COI haplotypes of *Hylurgus ligniperda* across all samples from native and non-native regions. Clades (including the two sub-clades 1a and 1b) were added based on those identified from the phylogenetic tree (Figure 2). The pies represent haplotypes, and sizes reflect the number of samples corresponding to haplotype sequences. Colours represent the country of origin of the sample.

### 3.4 Interception data

Our review of interceptions of *H. ligniperda* in Australia, Chile, Japan, New Zealand and the USA indicated that this species was encountered more than 800 times with imports into these countries alone (**Table 4**). The longest series of interception records of bark beetles (Scolytinae) exist for the USA (since 1914) and New Zealand (since 1950), whereas only more recent interceptions (since 1995, 2002, or 2003) were available for the other countries. In the USA, about 95% of interceptions originated from parts of the native range in Europe whereas in the other countries, interceptions from non-native ranges were predominant and accounted for 74 to > 99% of all interceptions.

**Table 4.** Number of interceptions of *Hylurgus ligniperda* recorded in Australia, Chile, Japan, New Zealand and the USA with known origins from native and non-native regions.

| Country of interception | Period of interception records | Europe (native) | North Africa (native) | West Asia (native) | Native Total | East Asia (non-native) | Oceania (non-native) | North America (non-native) | South America (non-native) | Southern Africa (non-native) | Non-native Total | Total known origins | % non-native out of known origins |
| --- | --- | --- | --- | --- | --- | --- | --- | --- | --- | --- | --- | --- | --- |
| Australia | 2003-2023 | 2 | 0 | 1 | 3 | 3 | 5 | 3 | 2 | 0 | 13 | 16 | 81.3% |
| Chile | 1995-2005 | 4 | 0 | 0 | 4 | 0 | 0 | 0 | 15 | 0 | 15 | 19 | 78.9% |
| Japan | 2002-2019 | 1 | 0 | 0 | 1 | 0 | 202 | 1 | 0 | 3 | 206 | 207 | 99.5% |
| New Zealand | 1950-2024 | 10 | 0 | 0 | 10 | 0 | 25 | 0 | 4 | 0 | 29 | 39 | 74.4% |
| USA | 1914-2008 | 515 | 0 | 3 | 518 | 8 | 6 | 0 | 10 | 4 | 28 | 546 | 5.1% |

Where longer-term data are available, changes over time in the number of interceptions of *H. ligniperda* were observed both in the USA and New Zealand (**Figure 4**). There were no interceptions of *H. ligniperda* from 1914 to 1963 and from 1950 to 1974 in the USA and New Zealand, respectively. Given that *H. ligniperda* has several easily recognised morphological characteristics and is common especially in southern Europe (Faccoli et al. 2020), the lack of reported interceptions probably means that it was not or only rarely moved with international shipments prior to the 1960s. However, since the 1960s or 1970s interceptions with imports originating from non-native ranges of *H. ligniperda* increased. For example, in New Zealand, interceptions from non-native ranges (in Oceania and South America) accounted for more than 80% of all interceptions of *H. ligniperda* between 1980 and 2009 (**Figure 4**). By contrast, in the USA, interceptions from non-native origins per decade reached a maximum of up to 6% of all interceptions in the 1970s and 1980s and declined afterwards. However, in both countries, interceptions with imports originating from non-native ranges coincided with prior establishments and growing populations of *H. ligniperda* in those regions (such as interceptions in New Zealand originating from other parts of Oceania and interceptions in the USA originating from Oceania and South America).

**Figure 4.**
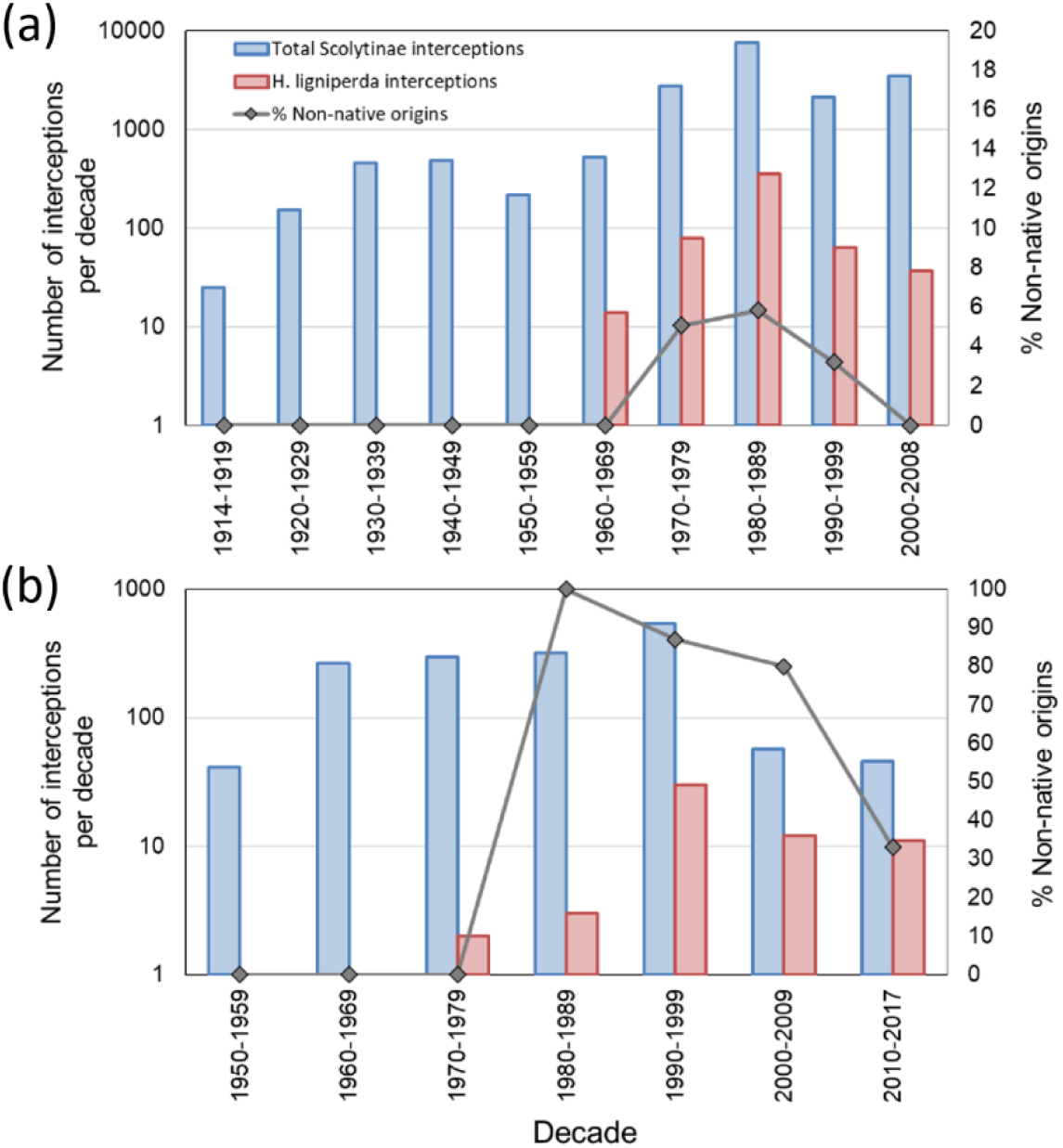
Interceptions at borders of (a) the USA and (b) New Zealand of all Scolytinae and of *Hylurgus ligniperda* only (primary y-axis), and the percentage of interceptions of *H. ligniperda* originating from non-native regions (i.e. origins where *H. ligniperda* occurs as a non-native species; secondary y-axis).

In the USA, more than 80% of interceptions of *H. ligniperda* were associated with wood packaging materials, especially wooden crates and dunnage, while for the remainder, in most cases it was not known or not recorded whether wood packaging was involved (**Table 5**). In New Zealand, wood packaging materials were also dominant, although there was a larger proportion where the association of wood packaging was not known or not recorded (**Table 5**). In terms of the commodities involved with these interceptions (which were recorded in detail only in the USA), about 60% of interceptions were stone and quarry products, especially tiles, granite, and marble (**Table 6**). Steel, machinery and parts accounted for nearly 14%, with the remainder being a wide range of other commodities as well as timber. Two cases of *H. ligniperda* as hitchhikers (e.g., beetles “at large” in shipping containers) were recorded.

**Table 5.** Wood packaging materials associated with interceptions of *Hylurgus ligniperda* recorded in the USA (1964-2008) and New Zealand (1975-2024) originating from native and non-native regions.

| Type of wood packaging | USA |  |  | New Zealand |  |  |
| --- | --- | --- | --- | --- | --- | --- |
|  | Native origins | Non-native origins | Unknown origins | Native origins | Non-native origins | Unknown origins |
| Crates (Casewood) | 315 | 15 | 1 | 1 | 6 | 1 |
| Dunnage | 100 | 7 | 0 | 3 | 7 | 1 |
| Pallets | 21 | 2 | 0 | 1 | 7 | 1 |
| No wood packaging | 6 | 3 | 1 | 0 | 1 | 0 |
| Unknown packaging | 75 | 0 |  | 5 | 8 | 22 |
| **Grand Total** | **517** | **27** | **2** | **10** | **29** | **25** |
| Percentage of all interceptions | 94.7% | 4.9% | 0.4% | 15.6% | 45.3% | 39.1% |

**Table 6.** Commodities associated with interceptions of *Hylurgus ligniperda* recorded in the USA between 1964 and 2008.

| Type of commodity | Europe | West Asia | Oceania | East Asia | South America | Southern Africa | Unknown | Grand Total |
| --- | --- | --- | --- | --- | --- | --- | --- | --- |
| **Stone** (Tiles, granite, marble, etc.) | 314 | 3 | 2 | 4 | 2 | 2 | 1 | **328** |
| **Steel** (Steel, machinery, parts, etc.) | 70 | 0 | 1 | 2 | 0 | 1 | 0 | **74** |
| **Other** (Glass, produce, etc.) | 56 | 0 | 3 | 2 | 0 | 1 | 0 | **62** |
| **Timber** | 4 | 0 | 0 | 0 | 3 | 0 | 1 | **8** |
| **Hitchhiker** | 2 | 0 | 0 | 0 | 0 | 0 | 0 | **2** |
| **Unknown commodity** | 68 | 0 | 0 | 0 | 4 | 0 | 0 | **72** |
| **Grand Total** | **514** | **3** | **6** | **8** | **9** | **4** | **2** | **546** |

## 4. Discussion

### 4.1 Native and non-native distribution

Our combined methods of (i) collecting specimens from all continents where host plants occur and (ii) a comprehensive review of the literature (including original reports dating back to the mid-1800s) provided much improved clarity about the true extent of the native and non-native ranges of *H. ligniperda*. We were able to correct the widespread misreporting of Madeira and the Azores as being part of the native range (e.g. Alonso-Zarazaga et al. 2023, CABI 2021, El Khoury et al. 2019, Wood and Bright 1992) when, in fact, neither *H. ligniperda* nor its host plants are native there (Crotch 1867, Méquignon 1942, Press 1994, Wollaston 1854). There are now 13 confirmed cases of independent invasions in geographically distinct regions, affecting all continents (except Antarctica), which makes *H. ligniperda* the most successful invader among true bark beetle species to date. Our synthesis of the current native and non-native ranges of *H. ligniperda* is presented in **Figure 5**.

**Figure 5.**
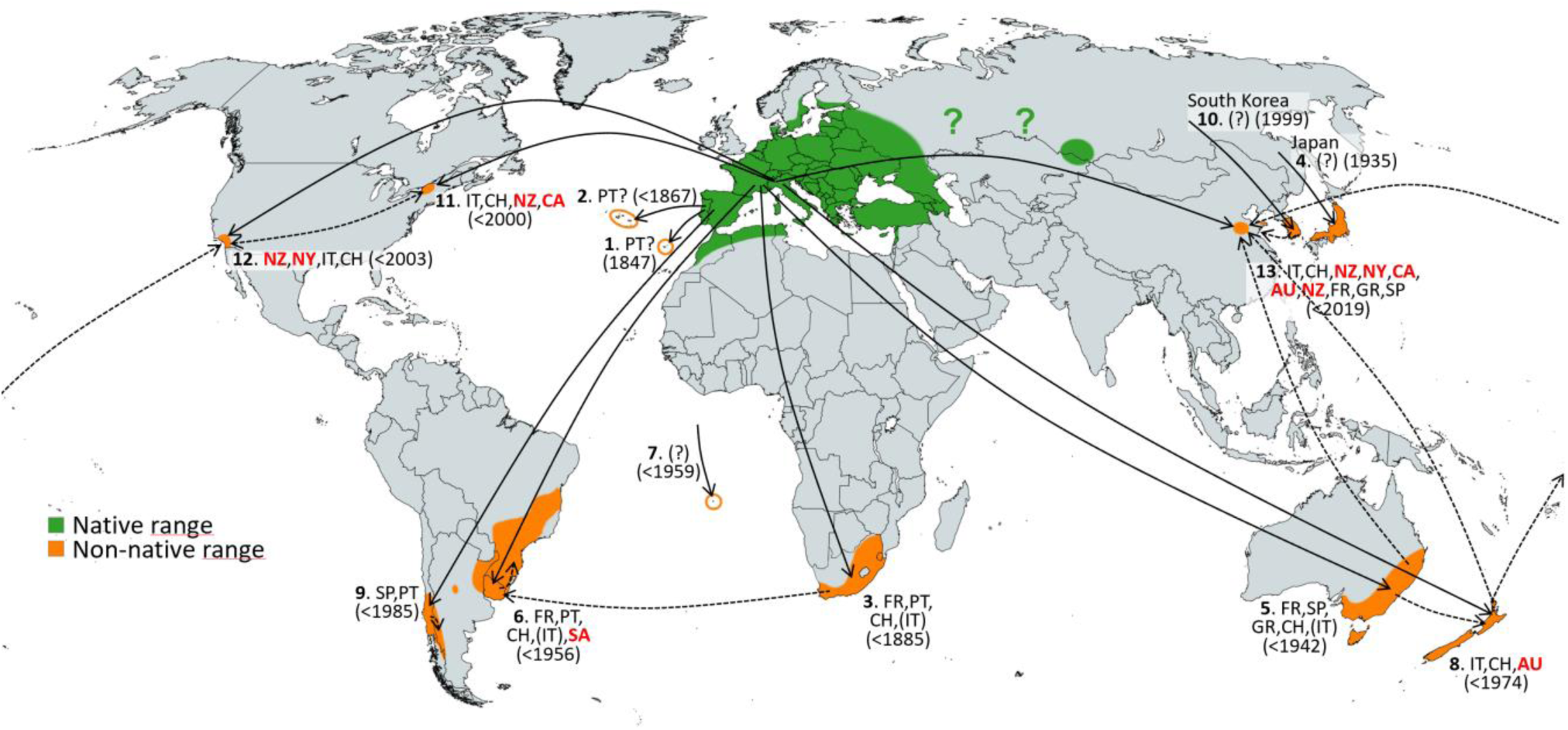
Reconstruction of most probable **invasion pathways** of *Hylurgus ligniperda* based on haplotype occurrences, temporal invasion sequences, and information on origins of imports with which specimens were intercepted. The numbers reflect the chronology of invasion events beginning with 1, Madeira (detected in 1847); 2, Azores; 3, South Africa; 4, Japan; 5, Australia; 6, Uruguay, 7, Saint Helena; 8, New Zealand; 9, Chile; 10, South Korea; 11, New York State (USA); 12, California (USA); 13, China. Two-letter codes are countries where specific haplotypes of non-native populations occur, representing potential source regions (AU, Australia; CA, California (USA); CH, Switzerland; FR, France; GR, Greece; IT, Italy; NY, New York State (USA); NZ, New Zealand; PT, Portugal; SA, South Africa; SP, Spain). Solid lines represent origins in the native range while broken lines and country codes in red font represent potential bridgehead invasions from non-native regions. Especially for more recent invasions of widespread haplotypes, multiple native and non-native origins are possible.

Some uncertainty about the distribution of *H. ligniperda* remains. While it has been clarified recently that it is not present in the Russian Far East or northeast China (Lin et al. 2021), its occurrence in western Siberia (Altai Krai) and northeast Kazakhstan appears to be well-documented (Kostin 1973, Stark 1955, Yanovsky 1999). Known records from the European part of Russia are limited to western regions, leaving an area north of Kazakhstan stretching about 2000 km west to east without any records of *H. ligniperda* (see “?” in **Figure 5**). This area should be climatically suitable for *H. ligniperda* as it is within the Köppen climatic region Dfb (humid continental climate) which also occurs in much of central and eastern Europe where *H. ligniperda* is found. Furthermore, pines (*Pinus* spp.) occur throughout this region, and there is a continuous presence of pines from northern Europe across northern Asia north of 50-55 degrees latitude (Nobis et al. 2012). Further studies would be necessary to clarify whether *H. ligniperda* is present in the area north of Kazakhstan and whether it is native or non-native in western Siberia.

While *H. ligniperda* has already colonised much of the available habitat in the southern hemisphere where pines had been introduced and the climate is favourable, most of the suitable habitats in North America and northeast Asia remain to be invaded. This agrees to some extent with the potential distribution of *H. ligniperda* according to maximum entropy modelling (Wu et al. 2022) although their modelling was not based on the entire non-native range, which would possibly increase the potential range.

### 4.2 Haplotype diversity and distribution in native and non-native regions

Our collections from the native range of *H. ligniperda* in Europe provided a representative if not complete overview of the phylogenetic diversity (in terms of haplotypes). Specimens from each of the six clades were found in the native range, including representatives of each of the three clades found in the non-native range. Although differences in the number of haplotypes among countries within the native range are probably influenced by the uneven sampling (e.g., 2 haplotypes from 3 samples in France vs. 6 haplotypes from 48 samples in Italy) and may not represent a true difference in haplotype diversity between countries in the native range, overall, our samples appear to reflect most of the haplotype diversity in the native range. However, there is a possibility that the single isolated haplotypes ‘Italy 1’, ‘Italy 2’ and ‘Portugal 1’ are part of other, larger clades. These individuals could also be representatives of other populations that arrived with imports from other parts of the native range. Such a situation has been documented for another scolytine species (Ruzzier et al. 2023).

Our analysis of COI nucleotide sequences revealed relatively high haplotype diversity in the native range in Europe. Even though we were not able to sample across the entire native range, we identified 21 haplotypes in the native range. These were spread across six well-supported clades, all of which were found within the native range. Nine haplotypes in the native range were singletons, suggesting that there may be more undetected haplotypes. However, most haplotypes were restricted to only one or two countries and regions in the native range, which suggests that limited dispersal has occurred among native populations.

Non-native specimens from five continents represented only about half as many haplotypes (12) as the native ones even though our non-native samples accounted for nearly three times as many samples and were collected across a larger geographic range. Only three of the six clades occurred in the non-native range, and these were divided geographically with sub-clade 1a occurring only in South America and South Africa, sub-clade 1b in Oceania and China, clade 6 in South America, and the most widespread clade 2 in Oceania, the USA and China. Eight haplotypes were only found in the non-native but not the native range, three of which with higher frequencies (HL-H2, N=35 in AR + UR; HL-H7, N=6 in CL, and HL-H23, N=16 in AU). Given the relative stability of COI, as it is not affected by recombination and has a relatively slow mutation rate, it is possible that these are ‘ghost haplotypes’ that are also present in the native range but were not sampled by us. However, all these missing haplotypes were closely related to other haplotypes occurring within the same parts of the non-native range. Therefore, it cannot be ruled out that these missing haplotypes have arisen locally in the non-native population via random mutations. In contrast to the nucleotide diversity, the 29 haplotypes showed identical amino acid sequences in both native and non-native populations, indicating strong purifying selection pressure, consistent with mitochondrial genome analyses of invasive *H. ligniperda* in China (An et al. 2024). The occurrence of so-called transient polymorphism (in the absence of alterations to protein sequences) has been documented previously in the context of invasive insect species, and it is attributed to changes in selection pressure that may favour substitutions in the new habitat (D’Ercole et al. 2023).

As with the number of haplotypes and clades, haplotype diversity was significantly lower in the non-native range than in the native range. This is consistent with the commonly observed pattern of non-native populations having lower genetic diversity after passing through a genetic bottleneck associated with the establishment of a small founder population (Bierman et al. 2022, Bock et al. 2015, Heckwolf et al. 2021). Regarding bark beetles in particular, lower genetic diversity was observed for instance in non-native populations of *Dendroctonus valens* in China, compared with native populations in North America (Taerum et al. 2016). Such bottlenecks may have strong negative effects on invasive populations, for example, by reducing fitness or limiting adaptive evolution (Dlugosch and Parker 2008), although this does not always play out as expected (Estoup et al. 2016). Non-native populations can increase in genetic diversity due to further arrivals and admixture of conspecifics from other, genetically different, source populations (Dlugosch and Parker 2008) because they can breed with individuals of previously established populations. However, we did not observe this in *H. ligniperda* as even the longest established non-native population we assessed did not have more than two or three haplotypes per country. This lack of genetic diversity did not inhibit the apparent success of non-native populations and their remarkable population growth.

### 4.3 Reconstruction of invasions and plausible origins of populations

#### Madeira and Azores

The earliest establishments of *H. ligniperda* in Madeira (<1847) and the Azores (<1867) are likely to have originated from Portugal because the islands were mainly colonised by Portuguese settlers who also established plantings of maritime pine (*Pinus pinaster*) in the nineteenth century (Pupo-Correia 2015). However, we have no genetic evidence from specimens from Madeira and the Azores.

#### South Africa

The next recorded establishment was in South Africa (<1885). Almost all specimens we obtained from South Africa belonged to haplotype HL-H5 (in clade 1a) which occurs in France, Portugal, southern Switzerland and probably also in northern Italy (**Table 3**). Thus, the South African population most likely originated from this part of Europe. One specimen from South Africa belonged to HL-H26 which is also in clade 1a.

#### Japan

With the detection of specimens of *H. ligniperda* in 1935, Japan was also among the earliest establishments. Although we were not able to obtain DNA of specimens from Japan, it is likely that the population originates from an introduction from the native range because at that time, non-native populations elsewhere were still limited and not highly abundant. However, more recently, numerous interceptions were recorded with imports from Oceania (**Table 4**), so that it is possible that the haplotypes occurring there may also be present.

#### Australia

In Australia, where a population was detected in 1942, one of the two haplotypes occurring there (HL-H3 in clade 1b) is shared with France, Greece, Spain and southern Switzerland. As there were no prior invasions in areas where this haplotype occurs, it is highly likely that the origin of the Australian population originated from one of these countries (or from areas in northern Italy near the areas where our Swiss specimens were collected). Other specimens from Australia were of haplotypes HL-H23 and HL-H29 which are also in clade 1b and closely related to HL-H3.

#### Uruguay, Brazil and northeast Argentina

Uruguay was the first country in South America where an established population of *H. ligniperda* was detected in 1956. We detected three haplotypes in Uruguay, HL-H2 and HL-H27 (two closely related haplotypes in clade 6 which were not recorded in Europe, but which are close to HL-H20 from Italy), and HL-H5 which occurs in France, Portugal, southern Switzerland (and probably also in northern Italy), and in South Africa (**Table 3**). The populations of HL-H2 and HL-H27 probably originated from an unsampled population in Europe. Given that an established population of *H. ligniperda* has been present in South Africa at least since the late 1800s, it is possible that the population of HL-H5 in Uruguay originated either from southern Europe or, via a bridgehead invasion, from South Africa (**Figure 5**). It was probably this population in Uruguay which later expanded into southern Brazil (where it was detected in 1991) and northeastern Argentina, possibly via Brazil (**Figure 5**).

#### Saint Helena

We have no information about the origin of the population in Saint Helena in the southern Atlantic, where *H. ligniperda* became established before 1959.

#### New Zealand

In New Zealand, where *H. ligniperda* was detected in 1974, two haplotypes were found: HL-H3 which is shared with France, Greece, Spain, southern Switzerland, and Australia, and HL-H4 which is shared with northern Italy and southern Switzerland as well as several other non-native regions which were colonised later. As *H. ligniperda* was frequently intercepted with imports from other parts of Oceania, and more so than with imports from Europe, there is a possibility that HL-H3 originated from Oceania or Europe, or from both (**Table 4**). However, based on our information on HL-H4, New Zealand was the first country outside the native range where this haplotype was recorded, so that it originated probably from an area south of the European Alps (**Figure 5**).

#### Chile and southwest Argentina

A second invasion of *H. ligniperda* in South America occurred in Chile where a population separate from the one established in Uruguay was detected in 1985. Three closely related haplotypes in clade 1a were found in Chile, whereby HL-H1, which is shared with Portugal and Spain, was most common, while the unique HL-H8 and HL-H9 were less common (**Table 3**). Given that this was the first known non-native occurrence of HL-H1, it probably originated from Portugal or Spain (**Figure 5**). This Chilean population (of HL-H1) later spread across the Andes into Argentina where it was detected in 2007. We also found specimens of HL-H2 in Argentina which are likely to have originated from Uruguay where this haplotype is common, or from adjacent states of Brazil.

#### South Korea

Although we have not examined specimens from South Korea, where *H. ligniperda* was detected in 1999, Lin et al. (2021) report that Korean specimens have the same COI sequences as those from China and New Zealand which suggests that haplotypes HL-H3 or HL-H4 or both are present.

#### USA (New York State and California)

In the USA, the two separate establishments in northern New York State and in southern California, detected in 2000 in 2003, respectively, are both of haplotype HL-H4, while in New York State we also found one specimen of HL-H28 (which is also in clade 2 and close to HL-H4) (**Table 3**). Although the two populations in the USA are separated by more than 3000 km, it is not clear whether they are entirely independent, given that they share the same haplotype. However, since both populations were still relatively small in the early 2000s, it is more likely that both were introduced from abroad. At the time of these establishments, probably in the 1990s (because of the time lag between establishment and detection), haplotype HL-H4 occurred in at least three regions, in southern Europe, New Zealand, and apparently also in South Korea. These could be the possible origin(s) of the populations in the USA (**Figure 5**). *Hylurgus ligniperda* was intercepted many times in the USA, mainly with imports from Europe, although some interceptions from East Asia, Oceania, South America and southern Africa have also been recorded (**Table 4**). Given the apparently more numerous arrivals from Europe, that is perhaps the more likely origin of populations in the USA.

#### China

The most recent documented establishments of *H. ligniperda* occurred in China. These were in Yantai and Tai’an (both in Shandong Province), first detected in Tai’an in 2019 (Lin et al. 2021), we found two haplotypes. Out of 23 sequenced specimens from Tai’an, six were of HL-H3 and 17 of HL-H4, while in Yantai, 25 out of 25 sequenced specimens were of HL-H4. This difference in composition of haplotypes was significant (Fisher’s exact test, P = 0.008) which suggests that two separate establishments may have occurred in Tai’an while there may have been only one establishment in Yantai. However, multiple establishments of each haplotype are also possible, as is the occurrence of a single mixed-haplotype establishment in Tai’an. Although we were unable to access comprehensive interception data from China, based on summaries by others, interceptions of *H. ligniperda* in China have been recorded with imports from “Australia, Uruguay and other countries” (Song et al. 2018 as cited by Yu et al. 2019) as well as New Zealand (Wang et al. 2019). Du et al. (2023) reported that live insects intercepted with wood packaging entering China originated from countries across all continents, especially from Africa, Europe and Southeast Asia, although they did not provide data on origins of *H. ligniperda* specifically. Given the widespread occurrence by the 2010s of both HL-H3 and HL-H4 on several continents, multiple origins from native and non-native origins are possible (**Figure 5**). The population of haplotype HL-H3 could have originated either from southern Europe, where this haplotype is widespread, or from Australia and New Zealand where this haplotype is present (**Table 4**). Populations of HL-H4 could have originated from southern Europe or from non-native ranges in New Zealand, South Korea or North America (**Table 4**). South America and South Africa can be ruled out as origins as neither HL-H3 nor HL-H4 have been found there.

### 4.4 Invasiveness and impacts of *Hylurgus ligniperda*

As a ‘secondary’ bark beetle which develops in the bark of recently died or felled trees at the base of tree stems, stumps, upper roots and logs in ground contact, *H. ligniperda* is not considered a pest in its native range (Eichhoff 1881, Fabre and Carle 1975). Eichhoff (1881) referred to its importance for forestry as “entirely irrelevant”. However, García de Viedma (1964) reported that in Spain, weakened pines up to 15 cm in diameter have been attacked and killed by *H. ligniperda* (as cited by Bain 1977). This appears to be the only such report from Spain (and Europe), and other forest entomologists in Spain consider *H. ligniperda* to be of no economic interest (Gil and Pajares 1986, M. J. Lombardero, pers. comm. 14 April 2025). *Hylurgus ligniperda* has generally received little attention from forest entomologists in Europe, and apart from Fabre and Carle (1975), few dedicated publications on its biology exist. This is despite its relative abundance, especially in southern Europe (Faccoli et al. 2020). Another indicator of its abundance in Europe is that it has been intercepted many times with imports from Europe as reported in the present study and by others (Brockerhoff et al. 2006a, Haack 2006). In fact, *H. ligniperda* is one of the most-intercepted bark beetles (Brockerhoff et al. 2006a, Haack 2006). Its association with pine timber (with bark) and wood packaging material, which is often made from pine timber (Krishnankutty et al. 2020) and, at least historically, often with sections of bark, have undoubtedly contributed to its movement with international trade and its numerous invasions. Both trade in timber and the widespread use of wood packaging materials (such as pallets, case wood and dunnage) with shipments of numerous commodities are recognised important pathways for the movement of bark beetles and other bark- and woodboring insects (Haack et al. 2014, Meurisse et al. 2019). In addition, *H. ligniperda* has been trapped in large numbers near ports in its native range (Rassati et al. 2014) which may also have contributed to its transport with international traded goods.

Apart from the frequent transport of *H. ligniperda* with international trade, the success as an invader has been facilitated by the widespread occurrence of host plants (*Pinus* spp.) across most temperate and many subtropical regions in the northern hemisphere (Nobis et al. 2012) and, in the form of plantations of non-native pines, also in large parts of the southern hemisphere (Lantschner et al. 2017). The presence of non-native plants has been identified as an important contributor to invasions of plant-feeding insects (Bertelsmeier et al. 2024). Furthermore, the spread of *H. ligniperda* has probably been enhanced further by its establishment in an increasing number of non-native ranges. In several cases, it has become highly abundant in its non-native range such as in Australia, New Zealand, and Uruguay (Brockerhoff et al. 2006b, Faccoli et al. 2020), and this is likely to have increased its propagule pressure via a bridgehead effect (Brockerhoff et al. 2025). The role of a bridgehead effect in invasions has been suspected or demonstrated in several cases involving insects (e.g., Bertelsmeier et al. 2018, Javal et al. 2019, Urvois et al. 2023). In the case of the wood wasp *Sirex noctilio*, which is also widespread as an invader and pine specialist, some populations were also suspected to have originated from non-native origins (Boissin et al. 2012). Another factor that may contribute to the invasiveness of *H. ligniperda* relates to its mating system. Most bark beetles have outbreeding mating systems (Kirkendall 1983) which create an impediment to successful establishment in a non-native area as mate finding is a prerequisite for successful reproduction, whereas species with compulsory or obligate inbreeding mating systems are expected to invade more easily as females are mated when they leave the gallery in which they developed. Although it is considered an outbreeding species, *H. ligniperda* may mate with siblings prior to host colonisation (Dacquin et al. 2025, Fabre and Carle 1975). This would also explain the relatively low Allee threshold and the small number of individuals needed for successful founder populations in experimental invasions (Chase et al. 2023). Furthermore, *H. ligniperda* is a capable disperser; it has been trapped up to 40 km from the nearest stands of host trees (Chase et al. 2017), and it responds strongly to host attractants (Kerr et al. 2017). Differences in invasiveness among haplotypes could also be involved. However, although haplotypes HL-H4 and HL-H3 are particularly common in invaded regions (with four and three invaded regions, respectively), we are not aware of any evidence that these haplotypes are inherently more successful as invaders than other haplotypes. Differences in relative propagule pressure from source regions is a more plausible reason for the apparently greater success of some haplotypes.

Contrary to its usually limited relevance in its native range, *H. ligniperda* has received much attention in invaded regions especially in the southern hemisphere but also in the USA and China. In Australia, it is generally of little concern, but outbreaks have been reported where planted *Pinus radiata* seedlings and even trees up to 14 years old were attacked and apparently even killed by *H. ligniperda* (Neumann 1987), although this seems to be happening very rarely if ever as a direct result of infestation by *H. ligniperda* in Australia. Reports from Chile suggest that *H. ligniperda* can attack and kill pine seedlings (Ciesla 1988), although Mausel et al. (2007) state that another non-native pine bark beetle, *Hylastes ater*, is mainly responsible for damage to seedlings. Ren et al. (2021) describe infestations of *Pinus thunbergii* by *H. ligniperda* with root damage occurring up to 1.5 metres below the soil surface. Another study in China (Bi et al. 2024) reports that “*H. ligniperda* and other boring insects can jointly harm the same pine tree, thereby accelerating the death of the tree” which implies that live trees were being attacked. These reports of damage to live seedlings and trees contrast with observations in New Zealand where forest health surveys and dedicated studies of attacks of pine seedlings and trees have never found *H. ligniperda* to be the responsible agent (Stephanie Sopow, pers. comm. 2025). Instead, attacks of live pine seedlings were exclusively caused by *Hylastes ater* (Reay et al. 2012, Sopow et al. 2015, Stephanie Sopow, pers. comm. 2025). Likewise, in a report on *Hylastes angustatus* and *H. ligniperda* in Eswatini, Bevan and Jones (1971) state “Unlike *Hylastes* it [*H. ligniperda*] has no obligatory maturation feeding on young green bark, and consequently it is not a pest of young living pines.” In New Zealand, damage or mortality of live pine trees have occurred, but this was caused by another insect, *Sirex noctilio* (Bain et al. 2012), or may have been due to drought or suppression by dominant trees. There are several potential reasons for the differences in impacts of *H. ligniperda* in different parts of its native and non-native ranges: (i) Misidentification of the agent actually causing death of seedlings or trees. It is possible that attacks by *H. ligniperda* of trees that are already dead are misinterpreted as the cause responsible for damage or mortality. (ii) The presence of other damaging insects or pathogens may pre-dispose trees and even kill them, leading to subsequent secondary attacks by *H. ligniperda*. In Australia, pines tend to be attacked by several other insects, including *Ips grandicollis*, *Sirex noctilio*, and *Essigella californica* (Nahrung et al. 2016), and more so than in New Zealand, which may explain different conclusions. (iii) Species are often more damaging in non-native ranges than in their native range, due to escape from natural enemies or new associations between pests and naïve host plants (Colautti et al. 2004, Enders et al. 2020, Liu and Stiling 2006); however, this by itself does not explain the difference between New Zealand and other southern hemisphere countries where the same host tree (*P. radiata*) occurs. However, this may be the case in China if the native Chinese pine *P. thunbergii* is indeed more susceptible to attacks by this bark beetle (Ren et al. 2021). (iv) Climatic differences could play a role as parts of Australia are drought-prone, which may weaken trees and increase their susceptibility to beetle attacks (Carnegie et al. 2022). (v) Given that different haplotypes occur in some of the invaded regions, it cannot be ruled out that populations in some regions are more damaging than in others. However, the same haplotypes occur in New Zealand, China and parts of the USA, which suggests that variation in reported damage are not related to different beetle genotypes.

While the role of *H. ligniperda* as a primary damaging agent is uncertain, its involvement as a vector of fungi causing sapstain on cut logs has been confirmed (Kim et al. 2011, McCarthy et al. 2013, Reay et al. 2006, Romón et al. 2014, Zhou et al. 2002). There are also concerns that *H. ligniperda* could vector tree pathogens such as *Leptographium wageneri*, the causal agent of blackstain root disease of conifers in western North America (Kim et al. 2011, Seybold et al. 2016), although it is unclear whether it could transmit the pathogen to live pine trees if it does in fact attack live trees. Irrespective of this uncertainty about a causal agent of tree damage, a main concern regarding *H. ligniperda* is its role as a ‘quarantine pest’ due to its occasional occurrence on export logs (Ren et al. 2021) which is unwanted by importing countries and necessitates quarantine treatments (van Haandel et al. 2017).

### 4.5 Conclusions

Our study provides an important update on the native and non-native ranges of *H. ligniperda*, one of the most successful invaders among forest insects. We determined that there were at least 13 invasion events of geographically separate regions across all continents. We corrected information from several erroneous references regarding native and non-native regions, some of which had been perpetuated in the literature for decades. Sequences of the COI gene from 464 specimens from eight native and nine non-native regions identified 29 haplotypes and showed lower genetic diversity in invaded than in native regions, typical of a genetic bottleneck in invading populations. Haplotypes of non-native populations showed clear spatial clustering indicative of independent invasions of different parts of the invaded ranges. Three haplotypes were shared between separate non-native regions, and especially HL-H4 was widespread across three non-native continents, suggesting the possible role of bridgehead invasions. This interpretation is also supported by numerous interception records originating from non-native regions, which shows the potential for non-native populations acting as origins of secondary invasions. However, as all the haplotypes that are most widespread and abundant in invaded regions were also more or less common in parts of the native range, and interceptions originating from these regions have been recorded, it cannot be determined with certainty what the original sources of invading populations were. Overall, our work is an important contribution to invasion biology which provides detailed insights into the roles of international trade and of widespread planting of non-native pines in invasions of *H. ligniperda* (and other host-specific pine-feeding insects) in regions which it would otherwise not have been able to colonise.

## Acknowledgements

We thank Don Booth (Bartlett Tree Experts, Charlotte, NC, USA), Georgia Dickson and Matt Scott (Scion, New Zealand), Simon Lawson (Forest Research Institute, University of the Sunshine Coast, Queensland, Australia), Åke Lindelöw (SLU, Sweden), Robert J. Rabaglia (USDA Forest Service, Washington DC, USA), Marie-Anne Auger-Rozenberg, Alain Roques, and Géraldine Roux (INRAE, Orléans, France) for assistance with the collection of specimens. Beat Ruffner, Sven Ulrich and Robin Winiger (all WSL, Switzerland) assisted in the laboratory or with genetic analyses. We also thank Stephanie Sopow (formerly Scion, New Zealand) for information on the lack of damages associated with *H. ligniperda*, and Maria J. Lombardero for publications from Spain and translations to English. Barney Caton (USDA PPQ, Raleigh, NC, USA), Andrew Liebhold (Czech University of Life Sciences, Czechia, formerly USDA Forest Service), Helen Nahrung (University of the Sunshine Coast, Queensland, Australia), Michael Ormsby (MPI, Wellington, New Zealand), Rebecca Turner (Scion / New Zealand Forest Research Institute, Christchurch, New Zealand) and Takehiko Yamanaka (NARO, AFFRC, Tsukuba, Japan) kindly provided interception data or helped with their interpretation. This project was partly inspired and supported by the National Socio-Environmental Synthesis Center (SESYNC) under funding received from the US National Science Foundation (DBI-1639145). MK was supported by the Ministry of Agriculture of the Czech Republic (Institutional Support MZE-RO0123). NIK was supported by the Russian Science Foundation (grant No. 22-16-00075).

