## Supplementary material for "Worldwide spread of *Hylurgus ligniperda* (Coleoptera: Scolytinae), and the potential role of bridgehead invasions": Suppl. Table 2 and caption Suppl. Table 1

### SUPPLEMENTARY MATERIALS

**Suppl. Table 1:** Detailed information about the geographic origin of the 464 *Hylurgus ligniperda* specimens analysed in this study (separate '.csv' file)

**Suppl. Table 2:** Occurrence of H. ligniperda as a native or non-native species by continent and country or region (see below)


Eckehard G. Brockerhoff[1], Lea Schläfli[1,2,3], Carolina Cornejo[1], Julia Kappeler[1,4], Jana Orbach[1], Amira Tiefenbacher[1,5], Quirin Kupper[1], Dimitrios Avtzis[6], Manuela Branco[7], Angus J. Carnegie[8], Kevin D. Chase[9], Juan Corley[10], Massimo Faccoli[11], Elizabeth Gilbride[12], Brett P. Hurley[13], Hervé Jactel[14], Jessica L. Kerr[15], Natalia I. Kirichenko[16], Miloš Knížek[17], Ferenc Lakatos[18], Victoria Lantschner[10], Gonzalo Martinez[19], Nicolas Meurisse[20], Miguel Angel Poisson[21], Adrian Poloni[22], Davide Rassati[11], Josep M. Riba-Flinch[23], José P. Ribeiro-Correia[1], Juan Shi[24], David Smith[25], Liam Somers[26], Yuan Yuan[27], Simone Prospero[1]

1 Swiss Federal Research Institute WSL, Birmensdorf, Switzerland

Information about all authors is given in the main document

**Suppl. Table 2** Occurrence of *H. ligniperda* as a native or non-native species by continent and country or region based on a major review of the literature, collection of specimens and correspondence with experts. Detailed information from key monographs is presented along with other key references as well as a synthesis of consolidated findings. The year of first detection or reporting as established in non-native regions is given. Regions of countries considered to be part of the native or non-native range are shown with green and orange highlighting, respectively.

| Continent | Region or country | Balachowsky (1949) | Browne (1968) | Wood and Bright (1992) | Pfeffer (1995) | Seybold et al. (2016) | Alonso-Zarazaga et al. (2023) | CABI (2021) | Lin et al. (2021). | Status (native/non-native) and year of detection in non-native countries | Key references and pers. comm. |
|---|---|---|---|---|---|---|---|---|---|---|---|
| **Europe** | | | | | | | | | | | |
| | **Western, Central, Northern and Eastern Europe** (incl. W Russia, i.e., west of the Ural Mountains) | Native (“Commun dans toute l’Europe tempérée …”) | Native (“Widely distributed in Europe”) | Native | Native (“Central Europe, Krim, Kaukasus”) | **Native** (but not present in Norway, and uncertain in Sweden (cite Schroeder 1990), and Estonia and Latvia (citing Bright and Skidmore 2002) | Native | Native | | **Native** (N France to Finland to Azerbaijan). Reported in [Great] Britain “but not as a breeding species” (Browne 1968, citing Duffy 1953). CABI (2021) incorrectly list it as ‘native’ in the United Kingdom (citing Wood and Bright 1992) (Duff 2018). No reports from Ireland. Absent in Norway (Kvamme and Lindelöw 2014, Torstein Kvamme, pers. comm. 18 May 2021). | Alonso-Zarazaga (2023, 2025), Seybold et al. (2016), Bright (2021) |
| | **Southern Europe** (incl. W Turkey) | Native (“Commun dans … certain points de l’Europe méridionale”) | Native (“Widely distributed in … the Mediterranean area”) | Native | Native (“Southern Europe”) | Native | Native | Native | | **Native** (Portugal to W Turkey). | Alonso-Zarazaga (2023, 2025) |
| | **Azores** | | | Native | | | Native | Native | | **Non-native** (**<1867**) “Species introduced with pine trees” Crotch (1867, p.362), “Importé aux Açores” (Méquignon (1942, p. 59). Note, pines are non-native in the Azores, neither is *H. ligniperda*. Year of first record unknown. | Crotch (1867), Méquignon (1942) |
| **Africa** | | | | | | | | | | | |
| | **North Africa** (Algeria, Morocco, Tunisia) | Not present (“Fait défaut en Afrique du Nord où il est remplacé par *H. micklitzi*”) | Native (“Widely distributed in … the Mediterranean area”) | Native (Morocco, Tunisia) | Native (Algeria) | Native | Native | Native (“Morocco and Tunisia”) | | **Native** (Morocco to Tunisia). | Alonso-Zarazaga (2023, 2025) |
| | **Canary Islands** | | Native (“Widely distributed in … the Atlantic islands”) | Native | | Native | Native | Native | | **Present and probably native but status uncertain**. Pines are native, but the islands are hundreds of km from the nearest populations of *H. ligniperda* on the mainland. | Crotch (1867) |
| | **Madeira** | Non-native (“introduit”) | Native (“Widely distributed in … the Atlantic islands”) | Native | | | Native | Native | | **Non-native** (detected ca. **1847**); earliest record of establishment given by Wollaston (1854). Also, pines are non-native in Madeira (Press 1994), so is *H. ligniperda*. | Wollaston (1854), Press (1994), Aguiar and Carvalho (2016), António F. Aguiar pers. comm. (12 February 2025) |

| | | | | | | | | | | | |
|---|---|---|---|---|---|---|---|---|---|---|---|
| | **South Africa** | | Non-native ("Parts of southern Africa" | Non-native | | Non-native | Non-native ("introduced in Afrotropical Region", no details) | Non-native | | **Non-native** (detected ca. **1885**). | Geertsema (1982), Schedl (1982), Tribe (1991) |
| | **Eswatini (Swaziland)** | | Non-native | | | Non-native | | Non-native | | **Non-native** (detected < 1971). | Bevan and Jones (1971) |
| | **Saint Helena** | | Non-native | Non-native | | Non-native | | Non-native | | **Non-native** (detected ca. **1959**, Year of detection unknown). | Schedl (1959) |
| **North America** | | | | | | | | | | | |
| | **USA** | | | | | Non-native | Non-native ("introduced in Nearctic Region", no details) | Non-native | | **Non-native** (**New York** detected in **2000**; **California** detected in **2003**; **Pennsylvania** detected in **2019**). | Hoebeke (2001), Haack (2006), Liu et al. (2007), Seybold et al. (2016), Robert J. Rabaglia (pers. comm. 01 July 2020) |
| **South America** | | | "Parts of South America" (no details) | | | | | | | | |
| | **Argentina** | | | | | | Non-native * | | | **Non-native** (detected ca. **2007**, near border with Chile). Meanwhile also in NE Argentina. | Tiranti (2010), Lantschner et al. (2024) |
| | **Brazil** | | | Non-native | | Non-native | Non-native * | Non-native | | **Non-native** (detected ca. **1991** near border with Uruguay). | Wood (2007), Lantschner et al. (2024) |
| | **Chile** | | | Non-native | | Non-native | Non-native * (* "introduced in Neotropical Region", no details) | Non-native | | **Non-native** (detected ca. **1985**). | Ciesla (1988) |
| | **Paraguay** | | | | | | | | | **Presence in Paraguay uncertain and not confirmed** (Wood 2007), if present then non-native. | Wood (2007) |
| | **Uruguay** | | | Non-native | | Non-native | Non-native * | Non-native | | **Non-native** (detected ca. **1956**). | Schedl (1966) |
| **Asia** | | | | | | | | | | | |
| | **West Asia** (Armenia, Cyprus, Lebanon, East Turkey) | | | Native (Turkey) | Native ("Minor Asia") | | Native (Armenia, Cyprus, Lebanon, E. Turkey) | Native (Turkey); Lebanon; | | **Native** (Armenia, Cyprus, Lebanon, E. Turkey). | Alonso-Zarazaga et al. (2023, 2025), El Khoury et al. (2019) |
| | **China** | | | Native (Present in "Manchuria in China") | | **Non-native** | Non-native ("native" in 2017 catalogue but "invasive" in 2023 catalogue) | Native | Non-native | **Non-native** (in Shandong detected ca. **2019**), native occurrence in NE China is incorrect, not present in NE China ("Manchuria") (Lin et al. 2021). | Lin et al. (2021), Alonso-Zarazaga et al. (2023) |
| | **Japan** | | Non-native | Native | | **Non-native** | Non-native ("native" in 2017 catalogue but "invasive" in 2023 catalogue) | Non-native | Non-native | **Non-native** (detected ca. **1935**). | Murayama (1936), Lin et al. (2021), Alonso-Zarazaga et al. (2023) |
| | **Russia, W Siberia** | | | | | | Native | | | **Native** in SW Siberia (Altai Krai) (Stark 1955, Yanovsky 1999) and N Kazakhstan (Kostin 1973); occurrence further east unlikely (Lin et al. (2021). | Stark (1955), Kostin (1973), Yanovsky (1999), Lin et al. (2021) |
| | **Russia, E Siberia** | | | | | | | | | **Not present** in the Russian Far East | Lin et al. (2021), Alonso-Zarazaga et al. (2023) |
| | **South Korea** | | | | | | Non-native ("invasive" in 2023 catalogue) | | Non-native | **Non-native** (detected ca. **1999**). | Park et al. (2017), Lin et al. (2021), Alonso-Zarazaga et al. (2023) |
| | **Sri Lanka** | | Non-native (no source given) | | | **Non-native** | | Non-native | | **Presence uncertain and unlikely**. Planted (non-native) pines occur in Sri Lanka, but no primary reference confirming the presence of *H. ligniperda* is known. | Browne (1968), CABI (2021) |
| **Oceania** | | | | | | | | | | | |

| | | | | | | | | | | | |
|---|---|---|---|---|---|---|---|---|---|---|---|
| | **Australia** | | Non-native (New South Wales, South Australia) | Non-native | | **Non-native** | Non-native (“introduced in Australian Region”, no details) | Non-native | | **Non-native** (detected ca. **1942**), widespread from South Australia to southern Queensland, also in Tasmania. | Swan (1942), Neumann (1987), GBIF.org (2025a) |
| | **New Zealand** | | | Non-native | | **Non-native** | Non-native (“introduced in Australia Region”, no details) | Non-native | | **Non-native** (detected in **1974**). | Bain et al. (1977), Brockerhoff et al. (2023) |
